# Sequential And Directional Insulation By Conserved CTCF Sites Underlies The *Hox* Timer In Pseudo-Embryos

**DOI:** 10.1101/2022.08.29.505673

**Authors:** Hocine Rekaik, Lucille Lopez-Delisle, Aurélie Hintermann, Bénédicte Mascrez, Célia Bochaton, Denis Duboule

## Abstract

During development, *Hox* genes are activated in a time sequence following their relative positions on their clusters, leading to the proper identities of structures along the rostral to caudal axis. To understand the mechanism operating this *Hox* timer, we used ES-cells derived stembryos and show that the core of the process involves the start of transcription at the 3’ part of the cluster, following *Wnt* signaling, and the concomitant loading of cohesin complexes on the transcribed DNA segments, i.e., with an asymmetric distribution along the gene cluster. Chromatin extrusion then occurs with successively more posterior CTCF sites acting as transient insulators, thus generating a progressive time-delay in the activation of more 5’-located genes due to long-range contacts with a flanking TAD. Mutant stembryos support this model and reveal that the iterated presence of evolutionary conserved and regularly spaced intergenic CTCF sites control the precision and the pace of this temporal mechanism.

## INTRODUCTION

*Hox* genes are essential for the organization of the body plan during animal development. In mammals, the transcription of these genes is deployed during gastrulation, at a time when the embryo produces and organizes its major body axis (see e.g. (Deschamps and Duboule, 2017). By the end of gastrulation, indeed, the embryo displays the classical distribution of *Hox* genes mRNAs, with progressively overlapping domains. As a consequence, cells at various anterior-posterior (AP) body levels express distinct combinations of HOX proteins, which instruct cellular populations as to which kind of morphologies they should produce (Kessel and Gruss, 1991; Krumlauf, 1994).

In vertebrates, the transcriptional activation of any *Hox* gene is largely fixed by its relative position within its genomic cluster (Gaunt et al., 1988), an unusual property initially described in flies (Harding et al., 1985; Lewis, 1978) and observed in most animals displaying an antero-posterior axis (Duboule and Dolle, 1989; Garcia-Fernàndez and Holland, 1994; Graham et al., 1989). In vertebrates, this mechanism is linked to a time-sequence in the transcriptional activation, first observed in mammals (Dolle et al., 1989; Izpisua-Belmonte et al., 1991) and subsequently generalized (Durston et al., 2012; Gaunt, 2015) with some controverses (see (Durston, 2019; Kondo et al., 2019). While the function of this timer was discussed in many instances (see (Durston, 2019; Gaunt, 2018; Kmita and Duboule, 2003), its mechanism has remained unknown, mostly due to the difficulties to experiment with the few NMP cells which feed the elongating axis with new mesoderm and neurectoderm tissue (Tzouanacou et al., 2009; Wilson et al., 2009) and where *Hox* genes are activated during axial extension.

A model was initially proposed whereby a progressive and directional opening of a closed chromatin configuration would lead to a stepwise accessibility of neighboring genes to activating factors (Noordermeer et al., 2011; Soshnikova and Duboule, 2009), with the onset of activation depends on *Wnt* signaling (Neijts et al., 2016), a signaling system active at the most posterior part of the developing embryo (Neijts and Deschamps, 2017). In subsequent phases, *Cdx* transcription factors were reported to activate more centrally located *Hox* genes (Amin et al., 2016; Mazzoni et al., 2013; Neijts et al., 2017), while *Gdf11* signaling might regulate more 5’-located (posterior) genes (Aires et al., 2019; Gaunt et al., 2013).

Coincidentally, the mapping of both the locations and orientations of CTCF binding sites within *Hox* gene clusters revealed three main sub-domains; first an ‘anterior’ domain devoid of CTCF sites; then a more centrally located domain where series of CTCF sites mostly show an orientation towards the 3’ end of the clusters, and a posterior domain where several CTCF sites display the opposite orientation (Amândio et al., 2021). This organization of CTCF sites in three domains is highly conserved either between species (Yakushiji-Kaminatsui et al., 2018) or even between paralogous *Hox* gene clusters (Amândio et al., 2021), i.e. over several hundred millions years of evolution, raising the hypothesis that these sites may be instrumental in the time-sequenced activation of the interspersed *Hox* genes. Indeed, CTCF is known for its potential to organize chromatin structures through the making and stabilization of large loops in combination with the cohesin complex (Fudenberg et al., 2016; Sanborn et al., 2015).

Here we address this question by using pseudo-embryos referred to as gastruloids (Turner et al., 2017) derived from an aggregate of ES cells cultivated *in vitro* for several days. After activating *Wnt* signaling for 24h, such ‘stembryos’ (Veenvliet et al., 2021) start to elongate a protrusion that resembles the outgrowth and progression of the tail bud and where the *Hox* timer is implemented (Beccari et al., 2018). We looked at various indicators of transcription, at the dynamic of cohesin recruitment and accumulation and at the changing architecture of the *HoxD* locus in parallel with its activation, and show that the *Hox* timer starts with a *Wnt*-dependent activation of the CTCF-free part of the cluster, which triggers an increased loading of cohesin complexes over this transcribed region. This is rapidly followed by the step-wise transcriptional activation of genes in the CTCF-rich region after a 3’ to 5’ progression in loop extrusion, along with progressive changes in the chromatin architecture of the locus. We challenged this model by using mutant stembryos and further show that, while the first phase may be sufficient to introduce a 5’ to 3’ asymmetry in transcription, the highly conserved array of CTCF sites is used to organize and secure the sequence and the pace of this *Hox* timer critical for the proper morphological stability of our axial structures.

## RESULTS

### Time-course of *Hox* gene activation in stembryos

In gastrulating mouse embryos, *Wnt* signaling contributes to the formation of the primitive streak from epiblast cells, which thus allows the emergence of germ layers that will shape the elongating body axis. Likewise, in those stembryos cultured as initially described in (Beccari et al., 2018), a pulse of the *Wnt* agonist chiron (Chi) 48 hours (h) AA (after aggregation; thereafter ‘48h’) of murine ES cells (mES) i.e., between 48h and 72h, triggers the differentiation of these epiblast-like cells to form a multi-layered structure resembling the posterior part of the elongating body axis. We generated a 12h resolution time-course ChIP-sequencing (ChIP-seq) dataset for H3K27ac, a chromatin mark found at active enhancers and associated with transcription (Fig. 1A). We analyzed the various H3K27ac profiles over the *HoxD* locus and its two flanking regulatory landscape C-DOM and T-DOM, two TADs, the latter being split into two sub-TAD1 and 2 (see e.g. (Rodríguez-Carballo et al., 2020) (Fig. 1B, C).

**Figure 1.**
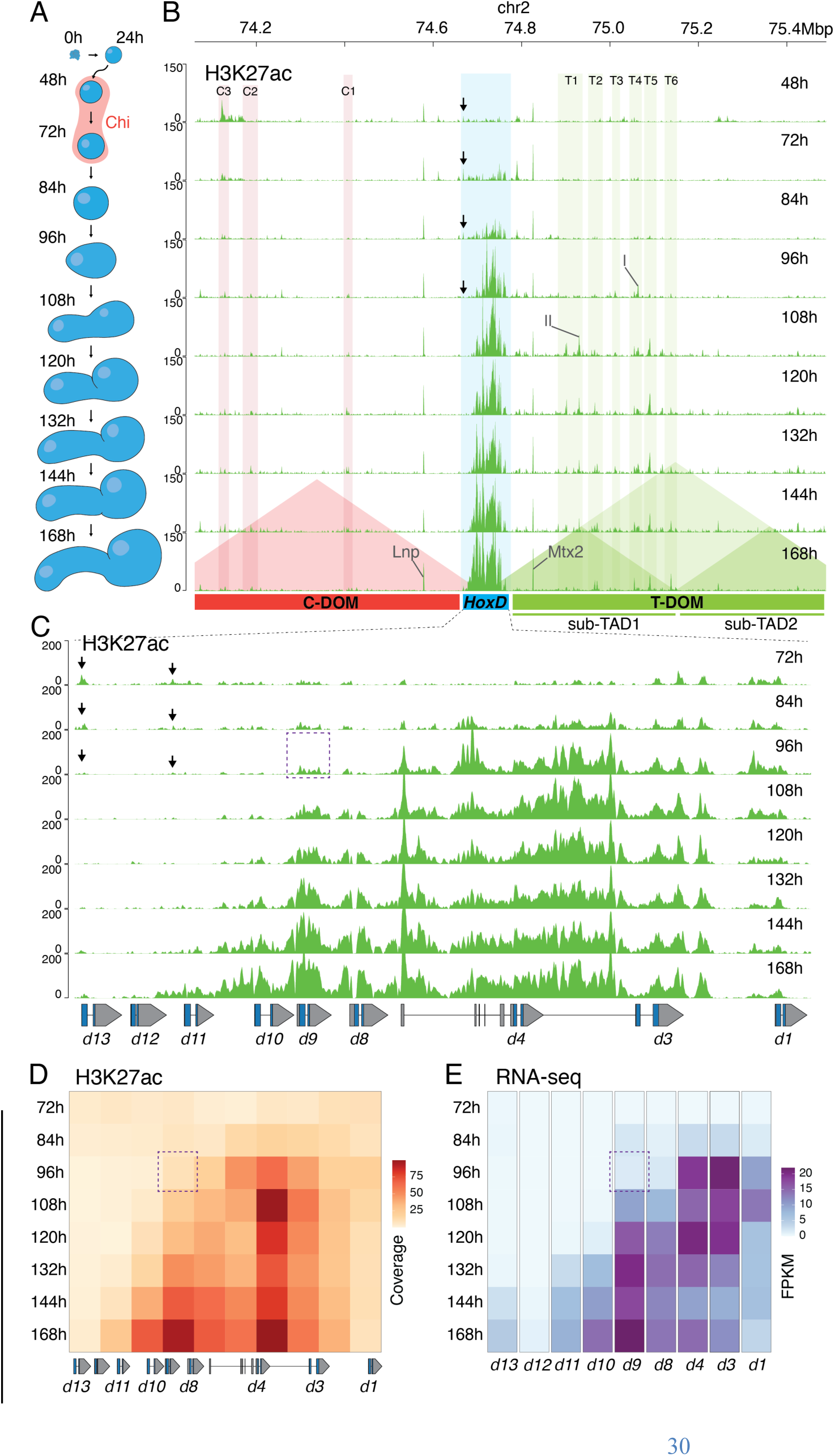
Sequential activation of the *HoxD* gene cluster upon *Wnt* signaling. (*A*) Various stembryonic stages used for the time course with the pulse of *Wnt* agonist in red. (*B*) Time-course of H3K27ac ChIP-seq over the entire *HoxD* locus. Genomic coordinates are on top and the positions of C-DOM and T-DOM are shown in red and green, respectively, at the bottom. The quantification of the H3K27ac signals within the colored vertical columns is in Fig. S1. Acetylation peaks at the *Mtx* and *Lnp* promoters are indicated. I and II point to the acetylation peaks used in Fig. S1. (*C*) Magnification of the *HoxD* cluster showing the progressive anterior to posterior spreading of H3K27ac. The start of robust coverage in the anterior part is counterbalanced by a decrease in the weaker acetylation initially detected in posterior regions (black arrows in *B* and *C*). (*D*) Heatmap of H3K27ac ChIP-seq coverage over the *HoxD* cluster at various time-points. Squares are 10kb large DNA segment. (*E*) Heatmap of FPKM values of *Hoxd* genes transcripts over-time (average of two replicates). The dashed region around *Hoxd9* at 96h in (*C*, *D* and *E*) highlights the increased activation in all subsequent stages, when compared to *Hoxd4*.

Soon after the end of Chiron treatment (72h), H3K27ac peaks appeared over the ‘anterior’ part of the cluster. From 72h to 84h, this signal covered mostly the *Hoxd1, Hoxd3* and *Hoxd4* genes, whereas no signal was detected in the flanking T-DOM. Acetylation signals started to appear within the T-DOM at 96h to become more and more prominent in the sub-TAD1, along with the increased acetylation of the *HoxD* cluster itself (Fig. 1B, 96h, green columns). Several enhancers controlling the expression of *Hoxd* genes in various tissues were already identified within sub-TAD1 (refs in (Amândio et al., 2021) and the dynamic of appearance of these peaks did not correlate with their distance to the *HoxD* cluster. Instead, they matched the time-window of the transcriptional activation of the cluster (Fig. S1). In contrast, no significant H3K27ac signal was scored in C-DOM (Figures 1B, pink columns; Fig. S1), further supporting the initial asymmetry in transcriptional regulation and the importance of T-DOM in this early activation.

Within the *HoxD* cluster, after the increase of H3K27ac marks over the *Hoxd1* and *Hoxd3* region at 72h, a rapid spreading of the acetylation marks was scored over the whole anterior part up to the border between *Hoxd4* and *Hoxd8* (Fig. 1C, D, 84h). At 96h, a slower and progressive acetylation of H3K27 was observed throughout the rest of the cluster, eventually covering the entire *Hoxd8* to *Hoxd13* segment at 168h (Fig. 1C, D). This multi-phasic dynamic, best visible through a time-lapse reconstruction (Supplementary Movie 1), was corroborated by a transcriptome analysis (Fig. 1E). *Hoxd1* to *Hoxd4* mRNAs were detected at 84h and became more abundant at 96h, a stage where *Hoxd8* and *Hoxd9* expression was low. The amounts of mRNAs from the latter two genes only increased at 108h. Likewise, expression of *Hoxd10* and *Hoxd11* was scored at 132h and *Hoxd13* at around 144h, correlating with the acetylation dynamic (Fig. 1D, E).

A low level of H3K27ac marks was scored over the entire posterior part of the cluster at 72h, which were erased right after the robust activation of the cluster at 96h (Fig. 1B, C, black arrows). Transcript from these genes were nevertheless not detected at these early time points, suggesting that the cluster was still in a bivalent chromatin configuration, as found in ES cells (Bernstein et al., 2005). Altogether, the initial transcription wave over *Hoxd1* to *Hoxd4* occurred before any T-DOM activity was scored, whereas a progressive activation of the rest of the cluster, starting at *Hoxd8* was clearly associated with a gain of H3K27 acetylation over many sequences in sub-TAD1, suggesting long-range regulation was at work to activate more central and posterior genes.

### Paused and elongating Pol II

To support these observations, we ChIPed-seq the large subunit of Pol II using a pan-specific antibody recognizing the C-terminal domain (CTD) of RPB1. Pol II occupancy was somewhat similar to the H3K27 acetylation profile, yet with an unexpected accumulation over *Hoxd8* and *Hoxd9* at 96h already (Fig. S2A), which was higher than both the H3K27ac profiles and transcripts levels (Fig. S2B, C; also see the boxed area in Fig. 1D-E). Consequently, while a colinear distribution was scored when comparing the 72h, 120h and 168h time points, profiles were virtually identical between 96h and 132h (Fig. S2A). This poor colinear progression was like that reported in amphibians (Kondo et al., 2019). However, when we ChIPed Pol II phosphorylated in serine 2 (Ser2-p) of the CTD domain, a modification that accompanies the transition from pausing to productive elongation, which usually covers the transcription unit with a robust enrichment in the 3’ part (e.g. (Zaborowska et al., 2016), a different distribution was scored (Fig. S2D).

The Pol II Ser2-p profile at 96h was clearly different from that of the pan-Pol II antibody and involved mainly the *Hoxd1* to *Hoxd4* region with a weak binding to *Hoxd8*, correlating well with the H3K27ac and RNA-seq profiles at this stage (Fig. S2D). The signals had now extended towards more posterior genes and was enriched in 3’of *Hoxd8* (Fig. S2D, left arrow in 84h), whereas at 144h, the signal had moved in 3’ of *Hoxd9* (Fig. S2D, arrowhead). At this latter stage, the coverage of the Pol II Ser2-p was eventually similar to that seen with the pan Pol II antibody. These differences between the two forms of Pol II (Fig. S2D, compare the signals with arrows in 96h to 144h) suggest that, while actively transcribed regions (Pol II Ser2-p positive) were spreading from the 3’ to the 5’ parts of the cluster, Pol II remained positioned throughout, likely under a paused state. Such striking differences were also readily observed at both *HoxA* and *HoxB* clusters (Fig. S3A, B, respectively). We concluded that elongating Pol II follows a colinear dynamic first detected in anterior regions of the clusters devoid of CTCF binding sites (see below), then progressively spreading towards posterior regions similar to H3K27 acetylation, unlike paused Pol II, which seems to be recruited in a rather simultaneous manner over an extended central part of the clusters including the transition from the 3’ CTCF-free region towards the CTCF rich region (Fig. 2A).

**Figure 2.**
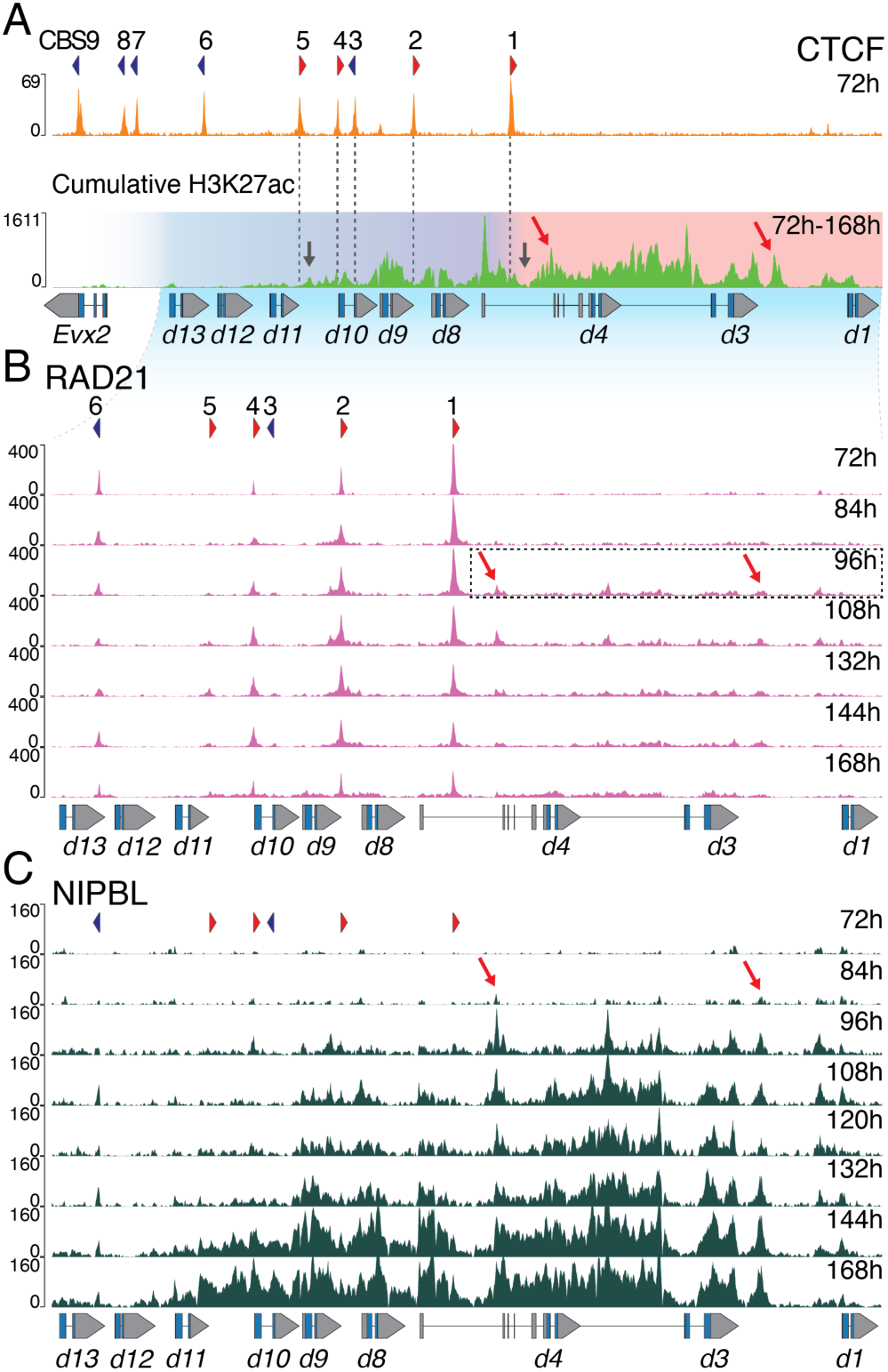
Dynamic and directional cohesin loading at the *HoxD* cluster. (*A*) CTCF ChIP-seq profile using stembryos at 72h (upper panel). The orientations of CBS1-9 motifs are shown with red and blue arrowheads. Below are cumulative H3K27ac ChIP-seq signals from 72h to 168h showing the various points of transitions in transcriptional activation over time. The red domain delineates the initial and rapid phase of acetylation, while the following and more progressive activation phase occurs in the blue domain. Black arrows point to regions where the progression in H3K27 acetylation was delayed (see Movie 1). (*B*) Time-course of RAD21 ChIP-seq profiles. CBS number and orientations are as in *A.* The accumulation of RAD21 outside CBSs is exemplified by the dashed box (quantifications in Fig. S5A). (*C*) Time-course of NIPBL ChIP-seq profiles over *Hoxd* genes. NIPBL binding corresponds to acetylated regions. Progressive NIPBL enrichment correlates with the activation dynamic of the cluster. Red arrows in *A, B* and *C* highlight two acetylated regions where NIPBL is detected and RAD21 accumulates.

### Distribution of bound CTCF in stembryos

This region of transition from a CTCF-free segment towards the first series of occupied CTCF sites (Fig. 2A, bottom; transition from red to magenta) was shown in mice to be important to set up proper expression domains in various embryonic tissues (Amândio et al., 2021; Narendra et al., 2016). We verified the binding profile of CTCF in stembryos and identified nine occupied CBSs inside *HoxD* (Fig. 2A; upper panel). Four out of the five ‘anterior’ CBSs displayed an orientation opposite to those CBSs located in the T-DOM (Figs. 2A, S5A; CBS1, 2, 4 and 5; red arrowheads), whereas the four CBSs located more ‘posteriorly’ showed an orientation toward CBSs located in C-DOM (Figs. 2A, S5A; CBS6, 7, 8 and 9, blue arrowheads). This pattern perfectly matched the profiles obtained with embryonic material (Amândio et al., 2021; Rodriguez-Carballo et al., 2017).

Since *Hoxd1*, *Hoxd3* and *Hoxd4* are activated concomitantly and their surrounding is devoid of any CBS, we asked whether the temporal progression observed in H3K27 acetylation may depend on the positioning of the CBSs. We produced a cumulative profile of H3K27ac signals between 72h and 168h, which showed transitions over time (Fig. 2A; lower panel). The CBSs systematically corresponded to poorly acetylated regions, some of them matching those places where the 3’ to 5’ progression in acetylation was slowed down, such as the posterior limit of the initial segment that was activated at ca. 84h and located close to CBS1, or the acetylation peak before *Hoxd11* facing CBS5, which remained unchanged until 132h (Fig. 2A, black arrows and Supplementary Movie 1). This suggested that the distribution of CBSs influences the progressive spreading of acetylation towards more posterior genes.

### Sequential and directional loading of the cohesin complex

We then assessed where cohesin complexes were recruited. We analyzed RAD21 binding, a component of this multi-protein complex. RAD21 was expectedly detected at several CBSs (Wendt et al., 2008), though with various enrichment levels (Fig. 2B, S4). Along with stembryonic development, low but significant levels of RAD21 accumulated outside CBSs, within gene bodies (Fig. 2B, black box). This accumulation of RAD21 over transcribed loci spread towards more posterior genes in subsequent stages, along with transcription (Fig. 2B, S4A compare 96h with 168h). To back up this dynamic of RAD21, we profiled NIPBL, one of the factors loading the cohesin complex onto chromatin (Ciosk et al., 2000)(Fig. 2C).

The NIPBL profile matched RAD21 binding in those regions outside CBSs, as illustrated by the particular enrichment over the *Hoxd1* to *Hoxd4* DNA segment between 84h and 96h, i.e., when RAD21 signals started to appear (Fig. 2B and C, between the red arrows). The NIPBL distribution then spread towards the posterior part of the cluster with a strong re-enforcement up to *Hoxd9* at 132h and reaching *Hoxd11* at 168h. NIPBL binding also followed the activation of the cluster, as evaluated by the acetylation dynamic (Fig. 1D) and NIPBL enriched peaks were all highly acetylated (Fig. 2A, lower panel; red arrows). Therefore, the progression of cohesin loading to actively transcribed region, which was quantified between each pair of CBSs (Fig. S4A) was likely triggered by NIPBL. Indeed, accumulation was enriched in the *Hoxd1* to CBS1 interval before it spread to CBS4 at 84h and reached the CBS4-CBS5 interval later at 168h, a stage where RAD21 was also detected throughout the *Hoxd9*-*Hoxd11* region. The profile of NIPBL at 168h precisely matched the H3K27ac profile, with signals covering *Hoxd11* and barely reaching the *Hoxd12* transcription unit. In many respects, these two profiles looked very similar, suggesting a tight link between the recruitment of NIPBL and transcription (Busslinger et al., 2017). NIPBL was slightly enriched at some promoters (Kagey et al., 2010; Zuin et al., 2014) and a massive coverage was scored over the entire transcribed domains.

### A self-propagating mechanism

We next analyzed the dynamic of RAD21 enrichment relative to CBSs orientations (Fig. S4B) and those with an orientation towards T-DOM (CBS1, 2, 4 and 5) showed variations in RAD21 accumulation reflecting their position within the cluster; RAD21 accumulation over CBS1 increased upon transcriptional activation of *Hoxd1* to *Hoxd4*, whereas the amount of CBS2, 4 and 5-associated RAD21 increased sequentially in time, following the progression of cohesin loading (Fig. S4B). We interpret this dynamic as the persistence at CTCF sites of cohesin complexes loaded more ‘anteriorly’ (in 3’) due to active transcription, which would extrude until reaching the next CBS with proper orientation. Consequently, the observed decrease in RAD21 accumulation at CBS1 reflects the extension of cohesin loading towards the posterior end of the cluster, leading to an averaging of RAD21 accumulation at all CBSs. This was supported by the absence of any consistent change in the accumulation of RAD21 at CBS6, 7, 8 and 9, which are oriented toward C-DOM, making them unable to accumulate cohesin complexes loaded onto the actively opening part of the cluster (Fig. S4B, bottom).

The relative accumulation of RAD21 also increased between 72h and 120h at some CBSs located within T-DOM (Fig. S5A, B; TD-CBS1, 2 and 5), which strongly interacted with the CBSs within *HoxD* due to their convergent orientations (Rodriguez-Carballo et al., 2017), in agreement with a function for these T-DOM CBS at these precise developmental stages (see below), for instance by bringing *Hoxd* genes to the vicinity of T-DOM-located enhancers. The extension of transcription towards a more ‘posterior’ part of the cluster would in turn recruit NIPBL and cohesin complexes, leading to some extrusion now starting also at more posterior positions thus leading to a self-propagating mechanism of successive transcriptional activation.

### Dynamic of Chromatin topology

We looked for modifications in chromatin structure during this dynamic process and produced a 24h time-series of Capture Hi-C (CHi-C) datasets, starting from 48h, when the activation of the cluster hasn’t yet started (Fig. 3A). At this stage, before *Wnt* signaling, interactions were observed between the *Hoxd* genes themselves, most likely as a result of their general coverage by polycomb marks (Noordermeer et al., 2011, 2014) leading to a ‘negative’ micro-TAD (Fig. 3A). At this stage, the cluster displayed several interactions with T-DOM, particularly with sub-TAD1 (Fig. 3A, 48h, arrows). Contacts established between CBSs in the cluster and sub-TAD1 were quantified (Fig. S6A, B, respectively) and CBS1 unambiguously established the strongest contacts when compared to other CBSs located within the *HoxD* cluster (Fig. 3A, white dashed line).

**Figure 3.**
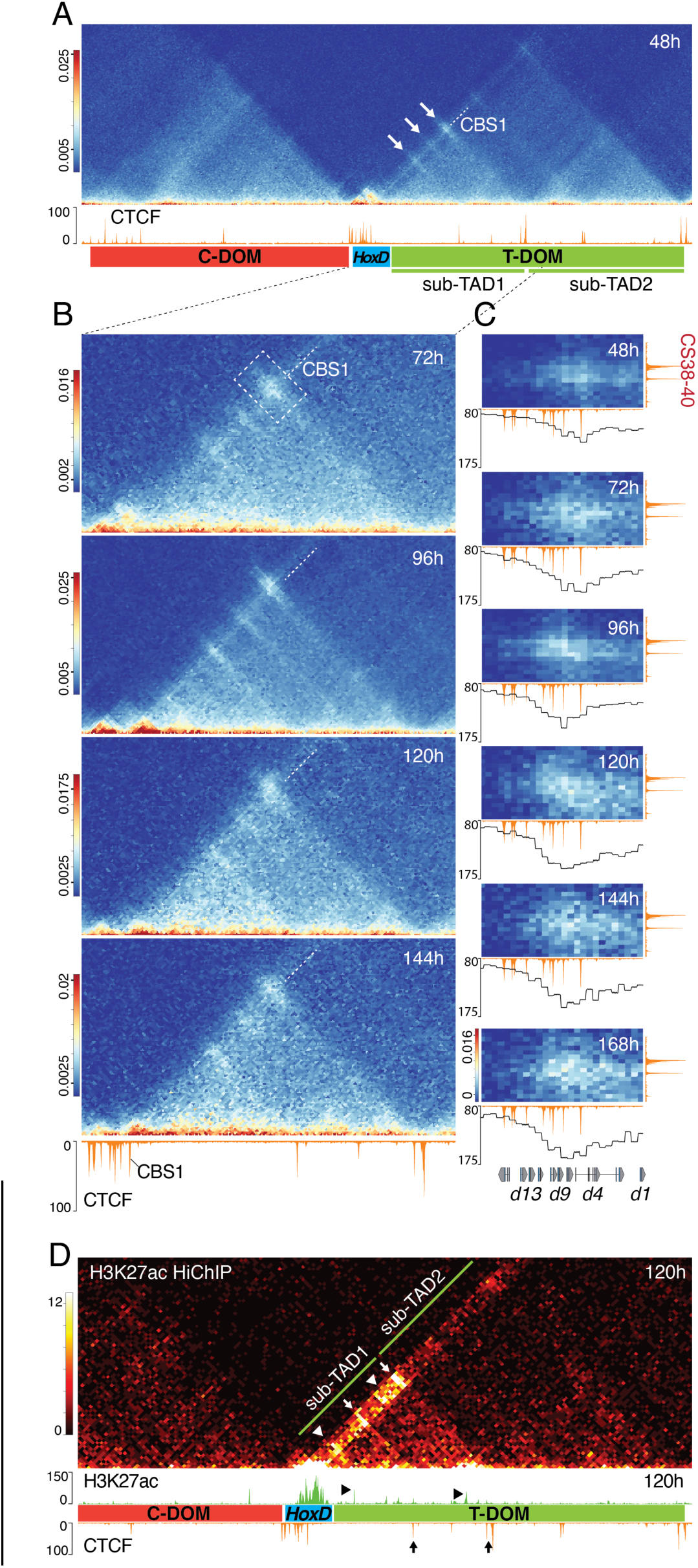
Topological reorganization upon transcriptional activation. (*A, B*) Time-course CHi-C at 5kb resolution using stembryos. (*A*) Contact maps including the *HoxD* cluster (blue rectangle) and both C-DOM (red) and T-DOM (green) TADs. A CTCF ChIP-seq profile is shown below to help visualize interactions. The arrows indicate strong initial contacts between CBS in the cluster and CBS within sub-TAD1. The line crossing all contact points and prolonged by a white dashed line shows contacts established by CBS1 (quantified in Fig. S7). (*B*) Magnifications of the *HoxD*-sub-TAD1 interval at various times points to show the transition of contacts initially established by CBS1 (dashed white line), then by more posterior CBS in the cluster (see the changing position of the strong contact points relative to the CBS1 dashed line (quantified in Fig. S7C)). The increase in contacts with sub-TAD1 also includes *Hoxd* genes body and were quantified (Fig. S7D, E). (*C*) Contacts heatmap between the gene cluster (x-axis) and the CBS-rich CS38-40 region (y-axis) at 48h to 168h. Bin size is 5kb. Below each heatmap is a CTCF ChIP track at 168h and a plot with the average value for each bin representing its interaction with the CS38-40 region. Interactions progressively involve more and more posterior CTCF sites. (*D*) H3K27ac HiChIP using 120h wild type stembryos (top), CTCF (168h) and H3K27ac (120h) (bottom). Bin size is 10kb. Higher contacts frequency is observed between the acetylated region in the *HoxD* cluster and sub-TAD1 when compared to sub-TAD2, with hotspots matching either the CBS or some acetylated peaks within sub-TAD1 (arrows and arrowheads, respectively).

At 96h, after *Wnt* activation, this distribution changed and the interactions between CBS1 and those CBSs associated with the CS38-40 region of sub-TAD1 (Fig. 3B, 96h; Fig. S6C) were no longer the most prominent, as interactions between this latter region and CBS2 became stronger (Fig. S6C). This shift in the balance of contacts towards the posterior part of the *HoxD* cluster paralleled the cohesin dynamic, which progressively engaged more posterior CBSs (see Fig. S4B), suggesting a link between the dynamic of cohesin loading and a topological reorganization of the locus. This was best visualized by fixing the position of contacts involving CBS1 with a white dashed line (Fig. 3B) and following the dynamic position of the main contact (Fig. 3B, boxed in 72h) moving up relative to this fixed line, i.e., involving now more posterior CBSs. This was supported by a quantification of contacts within this boxed area (Fig. 3C) with the curves below showing the distributions of contacts frequencies along with time, with a shift from the main use of CBS1 to the recruitment of more posterior CBSs (Fig. 3C, compare e.g., 48h with 120h). This shift can also be observed with the animation of this time-course dataset (Supplementary Movie 2).

These directional and stepwise increases in contact frequency with sub-TAD1 (or part thereof) also applied to the *Hoxd* genes bodies. *Hoxd1* to *Hoxd4* increased contact frequency with the whole sub-TAD1 at 96h (Fig. S6D). *Hoxd9* did not, but increased its interactions with the 3’ region of sub-TAD1 (Fig. S6E), a region rich in H3K27ac marks potentially reflecting enhancer-promoter contacts. These interactions were also documented by a HiChIP dataset for H3K27ac using 120h stembryos (Fig. 3D), a stage when anterior genes were activated and cohesin loading was already initiated. The contacts map produced a stripe illustrating the preferential interactions between the sub-TAD1 region and the active, H3K27ac-positive segment of the *HoxD* cluster, whereas contacts with sub-TAD2 were much less robust (Fig. 3D). The hotspots detected within the stripe matched either the sub-TAD1 CBSs (Fig. 3D, arrows), or acetylated peaks within sub-TAD1 (Fig. 3D, arrowheads), consistent with previous reports that had identified such stripes at active enhancers or within NIPBL-enriched regions (Kraft et al., 2019; Vian et al., 2018).

### Domain translocation and cohesin loading

To assess how these gains in contacts between sub-TAD1 and the *HoxD* cluster may relate to the initial cohesin loading region, we analyzed the CHi-C maps at a 2kb resolution, relative to the CTCF binding profile (Fig. S7A). At 48h, the negative micro-TAD was confirmed by a virtual 4C profile derived from the CHi-C experiment (Fig. S7B). Robust interactions were scored between a *Hoxd9* viewpoint and the entire cluster, from one extremity to the other (Fig. S7B, 48h, blue profile). However, a chromatin segment including CBS1, located between *Hoxd4* and *Hoxd8* showed much weaker interactions with the rest of the cluster. This translated into a ‘gap’ within the micro-TAD (Fig. S7A, top panel, double-arrows), also visible in the virtual 4C profile (Fig. S7B; double-arrows). This ca. 20kb large region thus looped out from the condensed structure, likely triggered by the interaction between CBS1 and the CBSs located within T-DOM (Fig. 3A, top; Fig. S7B, top).

At 72h, *Wnt*-dependent transcription of anterior *Hox* genes led to the end of contacts between the two halves of the gene cluster, as illustrated by the drastic translocation of the anterior part and the formation of two separate micro-TADs. This is best visible when comparing interaction maps of the gene cluster between 48h and 96h (Fig. S7A) as well as in the animation (Supplementary Movie 2). These two micro-TADs represent the two transcriptional states of each half of the cluster, inactive on the centromeric side (Fig. S7A, left) and active on the telomeric side (Fig. S7A, right), as shown by the mapping of H3K27me3 marks associated with silenced chromatin (Fig. S7C). The dynamic of intra-cluster interactions during this transition was also quantified (Fig. S8). Noteworthy, the H3K27me3 profile at 72h revealed a depleted region right around the position of CBS1, i.e., included into the fragment looping out of the negative ‘globule’ at 48h-72h (Fig. S7C, arrowhead). At 120h, H3K27me3 marks had been further erased towards the centromeric direction between CBS1 and CBS2, once again indicating the progression of transcription (Fig. S7C; vertical lines, arrow).

The H3K27ac and H3K27me3 profiles at 96h were mutually exclusive and corresponded to the two micro-TADs (schemes in Fig. 4A). The H3K27ac-positive domain (Fig. S7A, 96h, right) is that activated early on and where the recruitment of cohesin complexes was enriched. The H3K27me3-positive domain is the silenced part of *HoxD*. At 96h, the boundary between these two domains coincides with the position of CBS1 (Fig. S7C, arrowhead). Subsequently, the negative domain retracted until 120h when it became apparently stabilized (Fig. S7C; compare 120h with 144h), meanwhile the H3K27ac active domain expanded (Fig. S7A; 96h to 168h). The apparent stabilization of the H3K27me3 domain at late stages most likely reflected a dilution effect, for at these late stages, only those very posterior stembryonic cells activate new *Hoxd* genes and hence the vast majority of analyzed cells will keep their H3K27me3 domain unchanged. This progression was quantified by measuring, over-time, the chromatin interactions between the H3K27me3 domain borders in order to estimate its size (Fig. S8).

**Figure 4.**
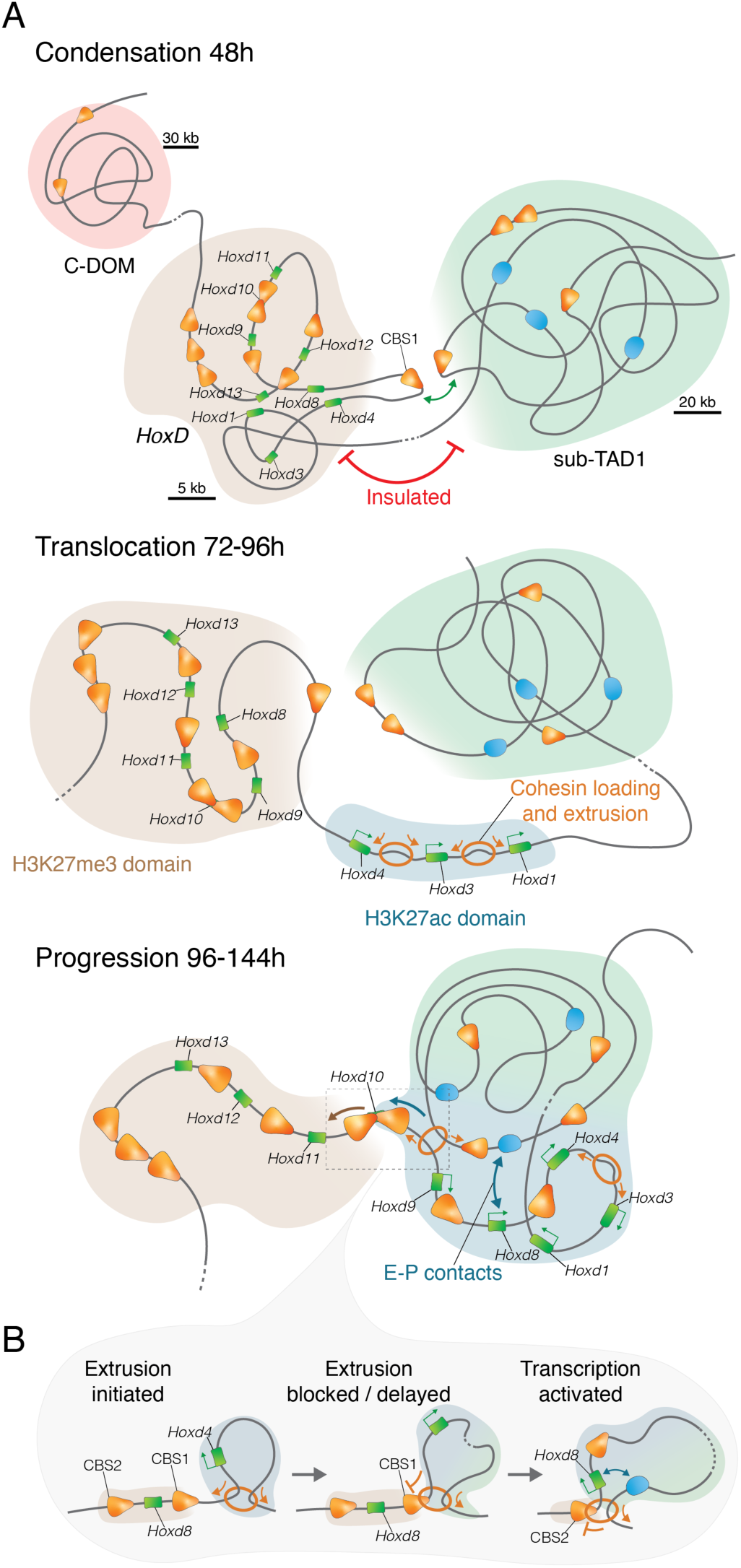
A model for the *Hox* timer. (*A*) Three phases in the dynamic activation of the *HoxD* locus. 1) Condensation: Initially, the cluster is in a condensed state (brown domain), covered by H3K27me3 marks. It is relatively well insulated from T-DOM and its sub-TAD1 (though with constitutive contact points), except for a ca. 20kb large DNA fragment containing CBS1, which loops out towards the sub-TAD1 located CBS (CBS1, green double arrow). 2) Translocation: While CBS1 keeps contacting the sub-TAD1 CBSs, *Wnt* -dependent transcriptional activation of anterior genes dissociates the cluster into two parts, with a now fully acetylated anterior domain where cohesin complexes will be preferentially loaded (blue domain). 3) Progression: Active loop extrusion in this transcribed part will initially reach CBS1 as a first block, to subsequently reach CBS2 with a given frequency, thus activating *Hoxd8* whose transcription will in turn recruit cohesin complexes and hence a self-entertained mechanism will spread towards the posterior part, concomitantly to a retraction of the H3K27me3 domain. (*B*) The ‘progression’ phase is tightly linked to extrusion by cohesin complexes, with an initial recruitment at anterior regions and a delayed progression caused by occupied CTCF sites of the proper orientations. CBS are represented by orange arrows indicating their orientation, genes are represented by green rectangles, enhancers by blue ovals and cohesin rings by orange circles.

When we considered interactions between the *Evx2* to *Hoxd12* border region and *Hoxd8*, we observed a decreased, already visible at 96h (Fig. S8, right). Contacts between *Hoxd9* and the *Evx2-Hoxd12* region decreased at 144h only, whereas the interactions with *Hoxd11* at 144h were only slightly decreased (Fig. S8, left). This progression occurred in parallel with the stepwise recruitment of novel CTCF sites in the cluster, which interacted with the main points of contact located within T-DOM (e.g., Fig. S5, TD-CBS1, TD-CBS2, TD-CBS5, Fig. S4B, 3A). In the Hi-C maps, this was illustrated by the transformation of discrete points of interactions into small bars of increasing length over time, the latter indicating several loops between one CTCF site in T-DOM and an increasing number of sites within the *HoxD* cluster (Fig. 3B, compare 72h and 144h, Supplementary Movie 2, white lines).

### A model for the *Hox* timer

These results suggest the following model (Fig. 4A). Initially, the cluster is packed into a negative H3K27me3 domain (Noordermeer et al., 2011; Soshnikova and Duboule, 2009) (Fig. 4A, top, brown domain). However, a central segment of the cluster containing CBS1 loops out and contacts several CTCF sites within the flanking T-DOM (Fig. 4A, green domain), in particular the CTCF sites present in the 38-40 region such as TD-CBS5 (Fig. 4A, top, contact between the brown and green domains). After *Wnt* signaling, *Hoxd1* is activated, rapidly followed by the entire anterior part of the cluster (*Hoxd3*-*Hoxd4*), i.e., up to CBS1. This activation coincides with a ‘translocation’ of this anterior part, as if the cluster was dissociated into two halves, one activated (Fig. 4A, middle, blue domain), the other one still silenced (Fig. 4A, middle, brown domain). Transcription in the active domain recruits cohesin complexes (Fig. 4A, middle, blue domain), which start to extrude DNA until extrusion reaches CBS1. Over time and at a given frequency (see e.g. (Gabriele et al., 2022), extrusion will bypass CBS1 and reach CBS2, which like CBS1 will also start looping towards the CBS sites located within sub-TAD1. The gene located in between will thus fall into the positive domain and become activated due to its proximity to T-DOM enhancers (Fig. 4A, bottom, E-P contacts). In turn, this newly transcribed gene will be targeted by NIPBL and cohesin will now be recruited more posteriorly, thus increasing the probability to bypass CBS2 and hit yet another CBS thus leading to a self-entertained mechanism progressing towards the posterior end of the gene cluster (Fig. 4B), after successive delays imposed by the presence of CTCF sites with an orientation towards sub-TAD1.

Surprisingly, this progression is not really visible with the ChIP-seq profile of Pol II, which at 96h is distributed even into the H3K27me3 domain (for example over *Hoxd10*), a profile that doesn’t evolve much during the next 48h. This overlap between Pol II and H3K27me3 is likely due to the heterogeneity of the process at the cellular level, depending of the AP position of cells with respect to the elongating stembryo, inducing a mixed interface between cells either positive or negative for any particular stage in the transcriptional progression. Also, the stable profile of Pol II (between 96h and 120h) reflects the presence of paused polymerases, as verified by the analysis of the phosphorylated Ser2 elongating polymerase, which in contrast showed a progression similar either to the epigenetic marks, or to the NIPBL profile. Following this model, after the *Wnt*-dependent initiation of the mechanism, the central part of the cluster (from the start of the CTCF binding sites (CBS1) to the TAD boundary between *Hoxd11* and *Hoxd13*) will be activated either through interactions with T-DOM enhancers, and/or to factors being able to access the cluster after its conformational modifications.

### Testing the model

To assess this model, we produced mutant stembryos out of a variety of modified ES cells carrying individual or multiple deletions of CBSs. The first mutant stembryos were deleted for CBS1 (Fig. 5A, *HoxD^Del(CBS1)-/-^* or Del(CBS1), as confirmed by ChIP-seq analysis (Fig. 5A). The normalized acetylation profile of Del(CBS1) mutant stembryos at 96h, when *Hoxd8* and *Hoxd9* transcription is still very weak in wild type, and at 120h when these latter two genes are now fully transcribed (Fig. S9B), showed enrichment over *Hoxd8* and *Hoxd9* at 96h and 120h. In contrast, the enrichment in the anterior part of the cluster was decreased in mutant stembryos (Fig. S9B). Inversion of enrichment ratio occurred right at the position of the deleted CTCF site and no gain in acetylation was observed for more posterior genes (e.g., *Hoxd10*) (Fig. S9B). These results suggested that a faster progression in the acetylation dynamic occurred in the cluster lacking the CBS1, this site acting as a barrier between the early activated *Hoxd1- Hoxd4* region and more centrally-located genes such as *Hoxd8* and *Hoxd9*. This was confirmed by RNA-seq at the same two time-points (Fig. 5B). Normalized expression level of *Hoxd* transcripts showed a decrease in expression of those genes located anterior to CBS1, whereas *Hoxd8* and *Hoxd9* mRNA levels were higher than in controls. A more robust effect was scored at 96h than at 120h for the gain of *Hoxd9* transcripts, while the *Hoxd8* mRNA level was back to normal at 120h, suggesting that CBS1 controls the timing of transcriptional activation of these genes rather than the maintenance of their transcription levels once activated.

**Figure 5.**
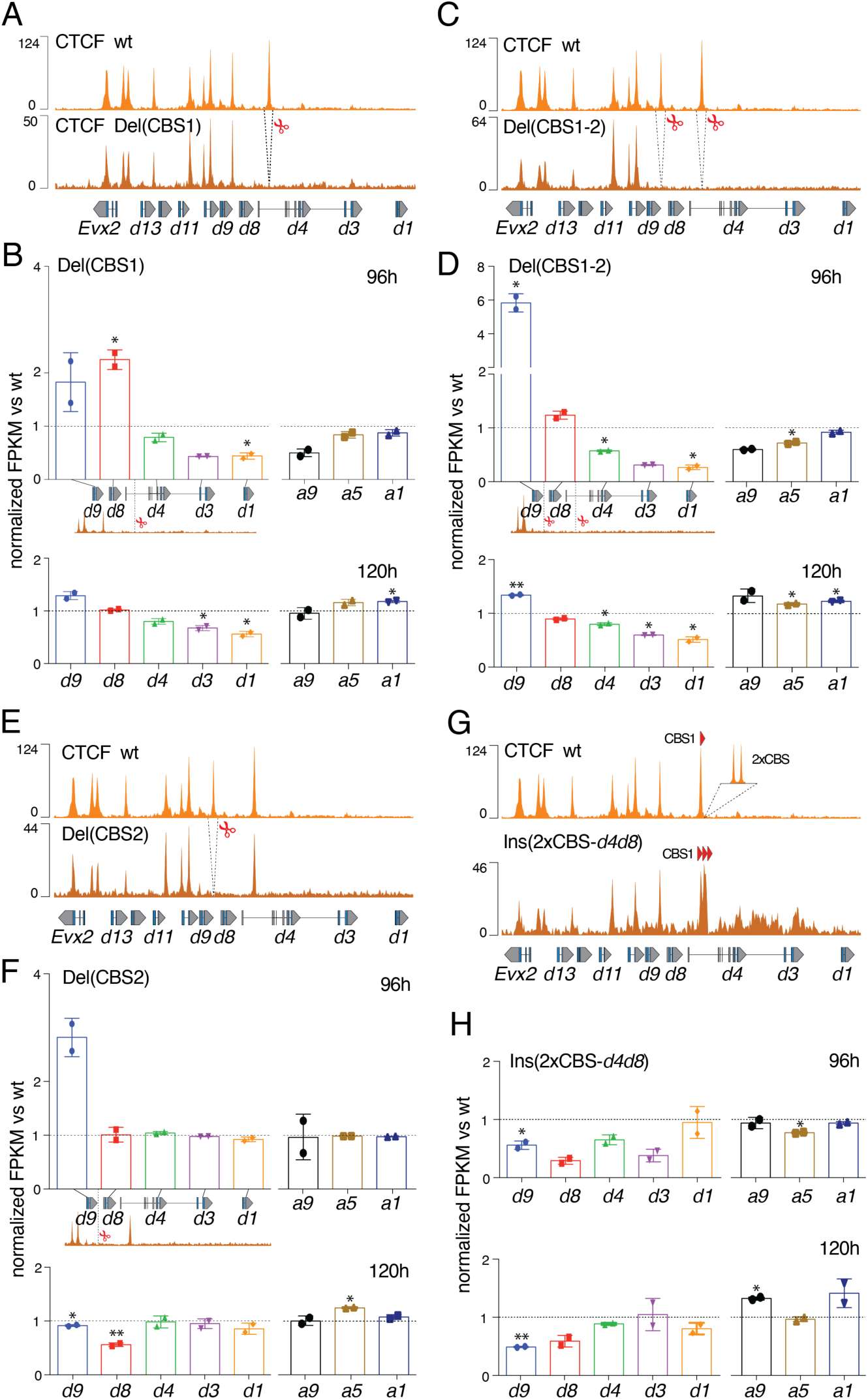
Deletions and insertion of CBS disturb the *Hox* timer. (*A*) CTCF ChIP-seq of either control stembryos at 168h (top) or Del(CBS1) stembryos at 96h (bottom). Scissors delineate the deletion. (*B*) Normalized FPKM values of *Hoxd* and *Hoxa* RNAs from Del(CBS1) stembryos compared to controls at 96h and 120h. The vertical line indicates the deleted CBS1. (*C*) CTCF control (top) and Del(CBS1-2) profiles at 96h (bottom). (*D*) Normalized FPKM values of *Hoxd* and *Hoxa* RNAs produced from Del(CBS1-2) stembryos and compared to controls at 96h (top) and 120h (bottom). (*E*) CTCF control (top) and Del(CBS2) profile at 96h (bottom). (*F*) Normalized FPKM values of *Hoxd* and *Hoxa* RNAs from Del(CBS2) stembryos compared to controls at 96h (top) and 120h (bottom). (*G*) CTCF control (top) and Ins(2xCBS-*d4d8*) profile at 144h (bottom). (*H*) Normalized FPKM values of *Hoxd* and *Hoxa* RNAs produced from Ins(2xCBS-*d4d8*) stembryos and compared to controls at 96h (top) and 120h (bottom). Values are represented as means ± SD. P-values were determined by Welch’s unequal variances t-test (* is p-value < 0.05, ** < 0.01). In (*E*), The number and orientations of CBS are indicated with red arrowheads.

We next analyzed stembryos carrying a double deletion of both CBS1 and CBS2 (Del(CBS1-2), a situation where the early transcribed genes and the more central part of the cluster were no longer separated by any occupied CTCF site (Fig. 5C). mRNA levels showed a gain in *Hoxd9* expression at 96h much higher than in the Del(CBS1)(Fig. 5D), in contrast with the moderate -if any-increase in *Hoxd8* transcripts level, which was below that observed when only the first CTCF was removed (Fig. 5B). There again, we observed a weakening of the effect of these two deletions at 120h, even though the tendency was clearly maintained (Fig. 5D, bottom). We concluded that the main upregulation effect when removing a CTCF site was observed for the gene located right in 5’ of the deleted site, i.e., *Hoxd8* for Del(CBS1) and *Hoxd9* for Del(CBS1-2), a conclusion further challenged by deleting CBS2 alone (Del(CBS2), Fig. 5E). In these mutants, expression of all anterior genes and of *Hoxd8* remained unchanged at 96h, whereas a clear increased in *Hoxd9* transcripts was scored (Fig. 5F), which again was no longer observed at 120h.

At this later stage, however, a decrease in the expression level of *Hoxd8* was observed, indicating that CBS2 may participate in the maintenance of *Hoxd8* expression. A similar effect was observed when CBS4 was deleted alone (Fig. S10A; Del(CBS4), since expression of *Hoxd8* and *Hoxd9,* the genes located anterior to the deleted CBS4, was decreased at 120h (Fig. S10B). In contrast, the *Hoxd11* mRNAs level, i.e., the gene located in 5’ of CBS4, was clearly upregulated at 144h. Overall, these results point to a dual function for these CTCF sites; on the one hand, CBSs are used as insulators for genes positioned posteriorly (in 5’), as exemplified by CBS2 and *Hoxd9.* On the other hand, CBSs are used as anchoring point for those genes located anterior (3’), as observed with CBS2 and *Hoxd8*. As a result, the deletion of a CBS site will tend to activate prematurely the gene positioned in 5’, while it will fail to maintain the expression of the 3’-located gene at its normal level.

### Insertion of supernumerary CTCF sites and interference with cohesin loading

We also inserted two supernumerary CBSs within a 2kb distance next to CBS1. In these stembryos, three CTCF sites instead of one are now ‘opening’ the series of CBSs, all oriented toward sub-TAD1 (Fig. 5G, Ins(2xCBS-*d4d8*). RNA-seq libraries revealed that the mRNA levels of several genes were significantly decreased at 96h (from *Hoxd3* to *Hoxd9*) (Fig. 5H, top). At 120h, the genes positioned posterior to the three compact CBSs (*Hoxd8* and *Hoxd9*) were still affected in their transcription, suggesting that the initial delay in expression of central genes was persistent in this case (Fig. 5H, bottom).

We also looked at the importance of loading cohesin in the anterior portion of the cluster by deleting this region. This deletion removed from *Hoxd1* to *Hoxd4* included, yet CBS1 was left in place (Fig. 6A). RNA-seq of mutant stembryos showed a marked decrease in transcripts level for *Hoxd8* and *Hoxd9*, both at 96h and 120h (Fig. 6B). To see whether this decrease in transcription was associated with a change in the contacts established between the various intra-cluster CBSs and sub-TAD1, we produced a CHi-C dataset of Del(*Hoxd1-Hoxd4*) stembryos at 96h (Fig. 6C). While the micro-TAD observed over the anterior region of the cluster under normal condition had expectedly disappeared along with the deletion (Fig. 6C, arrow; compare with Fig. S7A at 96h), a quantification of contacts between the various CBS1-5 and the CS38-40 sub-TAD1 region revealed important modifications. In 96h control stembryos, CBS2 showed a higher contact frequency with the CBSs located within the CS38-40 region, when compared to CBS1 (Fig. S11A). In contrast, the Del(*Hoxd1-Hoxd4*) mutant stembryos showed comparable interaction levels between CS38-40 and CBS1 and 2 (Fig. S11A). We interpret this as a reduction in cohesin complexes reaching more ‘posterior’ CBS, due to reduced loading in the absence of the DNA segment including early transcribed genes and targeted by NIPBL. Basal accumulation of cohesin was nevertheless sufficient to establish the expected constitutive contacts with the CS38-40 region (Fig. 6C, dashed box).

**Figure 6.**
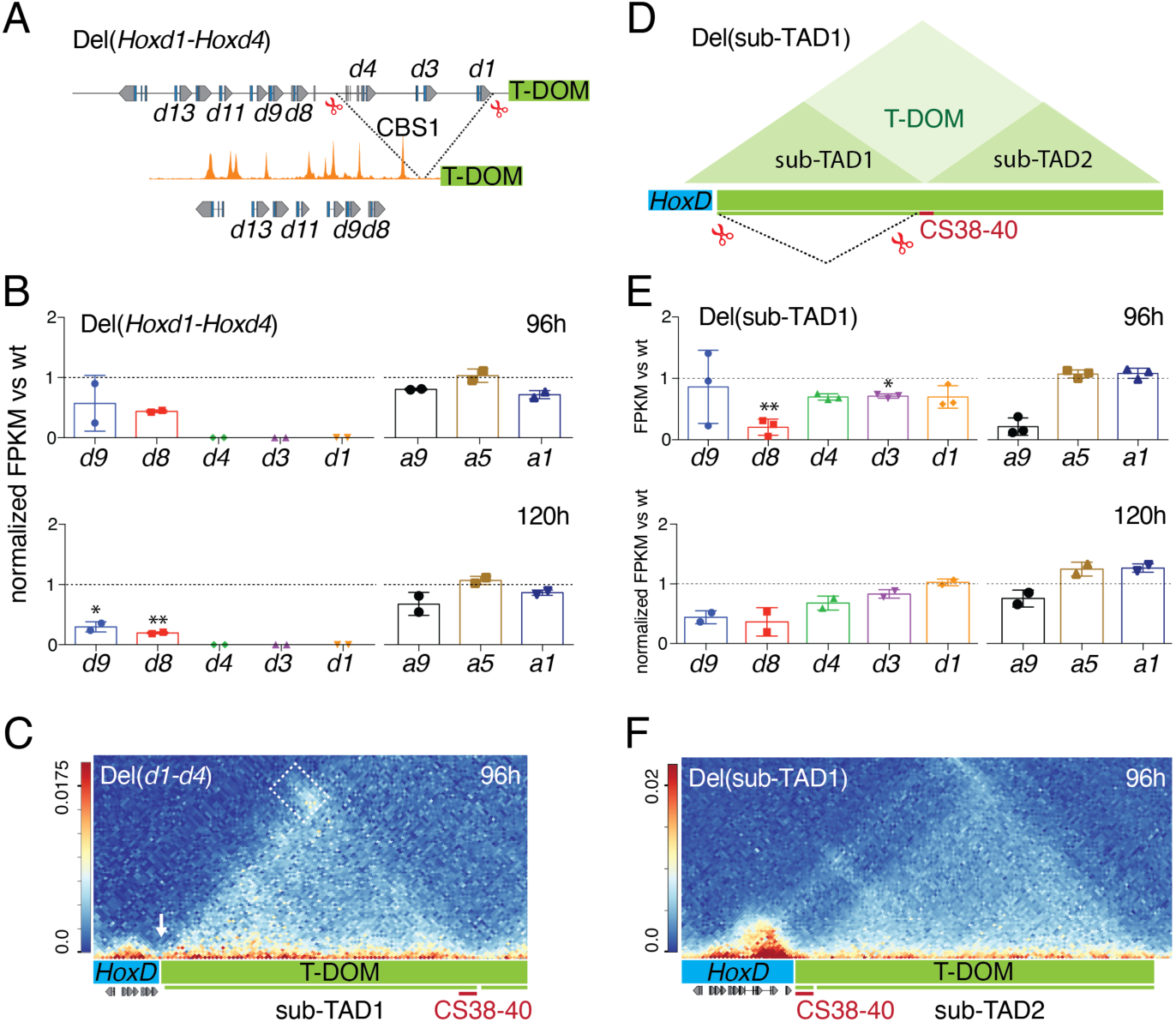
Larger deletions at the *HoxD* locus affect the timing of transcription. (*A*) The Del(*Hoxd1-Hoxd4*) mutant, with the anterior region of the *HoxD* cluster removed. (*B*) Normalized FPKM values of *Hoxd* and *Hoxa* RNAs from Del(*Hoxd1-Hoxd4*) stembryos compared to controls at 96h and 120h. (*C*) CHi-C map produced from Del(*Hoxd1-Hoxd4*) stembryos at 96h, mapped onto an *in silico* reconstructed mutant genome. Bin size 5kb. The deletion breakpoint is indicated with a white arrow. Contacts between the cluster and the CS38-40 region are scored (dashed box) and quantified in Fig. S11A. (*D*) Deletion of sub-TAD1. The boundary between sub-TAD1 and sub-TAD2 (CS38-40) is left in place. (*E*) FPKM (96h) and normalized FPKM (120h) values for *Hoxd* and *Hoxa* RNAs from Del(sub-TAD1) stembryos compared to control (96h: n=3 mutant *versus* 2 wt; 120h: n=2). Values are represented as means ± SD. P-values were determined by Welch’s unequal variances t-test (* is p-value < 0.05, ** < 0.01). (*F*) CHi-C map using Del(sub-TAD1) stembryos at 96h, mapped onto an *in silico* reconstructed mutant genome. Bin size 5kb. A higher contact frequency is observed between the *Hoxd* genes cluster and the CS38-40 boundary region, now relocated close by. Quantifications in Fig. S11B.

Next, we deleted sub-TAD1 to evaluate its importance for the transcription of those central genes located in 5’ of CBS1. Accordingly, the CBS-rich CS38-40 region was moved close to the anterior part of the cluster (Fig. 6D). Changes in mRNAs level were already observed at 96h for *Hoxd8* (Fig. 6E, top) and a substantial decrease was scored at 120h for both *Hoxd8* and *Hoxd9,* whereas anterior genes were less affected (Fig. 6E, bottom). These results confirmed that posterior *Hoxd* genes require sub-TAD1 to be properly activated and that this may be achieved through CBS-dependent interactions. It also confirmed that the initial *Wnt*-dependent activation of anterior genes is independent from sub-TAD1 and thus potentially involves elements located within the cluster itself. The CHi-C profile of these mutant stembryos revealed a rewiring of contacts between the cluster and the now closely located region CS38-40 (Fig. 6F), with a strong micro-TAD including the anterior portion of the cluster, where cohesin loading could presumably occur normally (Fig. S11B).

### CTCF as a time keeper

Lastly, we assessed the importance of CBSs in the transcription timing dynamic more globally by using stembryos lacking all CBS1 to CBS5 *in-cis*, as controlled by the absence of RAD21 accumulation up to CBS6, the CTCF site with an opposite orientation located between *Hoxd12* and *Hoxd13* (Fig. 7A). Expression of posterior genes was upregulated at all stages analyzed (Fig. 7B). At 120h, both *Hoxd10* and *Hoxd11* expression was gained as well as *Hoxd13* at 144h. Therefore, the absence of bound CTCF in the anterior and central parts of the cluster lead to the premature activation of those genes located 5’ to CBS1, supporting the view that loops stabilization by CTCF after extrusion may be key to regulate the pace of the transcription dynamics by retaining and focusing transcription for some time, before additional loading of cohesin and the extrusion mechanism goes through and reaches the next CBS.

**Figure 7.**
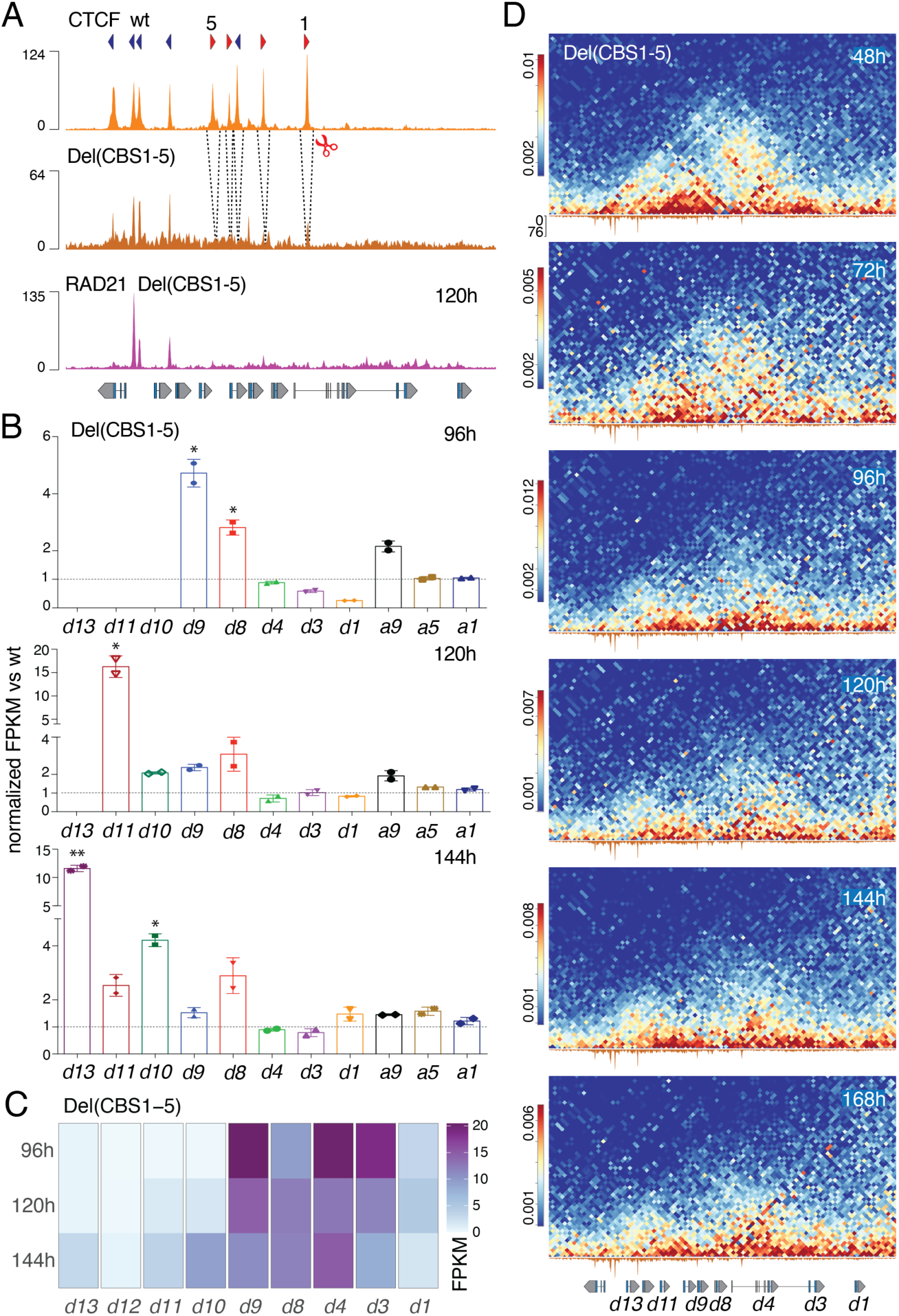
Impact of multiple CBS deletions *in-cis* on the *Hox* timer. (*A*) CTCF ChIP-seq data from control at 168h (top) and Del(CBS1-5) mutant stembryos at 96h (middle). Deleted CBS are indicated with dashed lines. The RAD21 profile at 120h (bottom) shows no accumulation at the deleted sites (compare with Fig. 2B). (*B*) Normalized FPKM values for *Hoxd* and *Hoxa* RNAs from Del(CBS1-5) stembryos compared to control at 96h, 120h and 144h. Expression of *Hoxd10* and *Hoxd11* was not considered at 96h, nor that of *Hoxd13* at 96h and 120h. Increased expression was observed for central and posterior *Hoxd* genes in mutant stembryos. Values are represented as means ± SD. P-values were determined by Welch’s unequal variances t-test (* is p-value < 0.05, ** < 0.01). (*C*) Heatmap of FPKM values for *Hoxd* genes transcripts in Del(CBS1-5) mutant stembryos at various time points (average of two replicates). (*D*) Time-course CHi-C using Del(CBS1-5) stembryos. Bin size is 2kb and libraries mapped onto wild type genome (mm10). When compared to control specimen (Fig. S7), the translocation (if any) of the cluster into two globules is very much perturbed.

Premature activation was confirmed by comparing normalized H3K27me3 profiles between control and CBS1-5 mutant stembryos at 96h (Fig. S12A). While the anterior part was mostly similar to control, the entire H3K27me3 domain up to the *Evx2* gene was ‘weakened’, with a particular loss over the *Hoxd11* to *Hoxd8* region (Fig. S12A, bracket in the subtraction profile at the bottom), i.e., the region containing CBS1 to 5. Therefore, in the absence of bound CTCF, the whole cluster became somehow transcriptionally leaky. However, the extent of the H3K27me3 domain did not change in mutant stembryos, indicating that bound CTCF do not interfere with the early distribution of polycomb marks over the gene cluster in ES cells. Despite this severe impact upon the timing of activation, a weak tendency to follow a colinear activation was still observed in particular at late stages and from *Hoxd9* onwards (Fig. 7C; compare 120h to 144h).

We produced a time-series of CHi-C datasets with Del(CBS1-5) stembryos and noticed several changes in the dynamic of chromatin architecture when compared to controls (Fig. 7D). First, the *Wnt*-dependent translocation of the gene cluster into two separated domains was no longer observed, supporting a requirement of CTCF for this process. Second, at 96h, the progression of contacts towards the more posterior (5’) part of the cluster was not observed in mutant stembryos, indicating again that it was normally driven by occupied CTCF sites. Instead, after activation of the *Wnt* pathway, the anterior part did not clearly segregate from the central and posterior regions of the cluster and contacts remained distributed throughout (Fig. 7D), reflecting a diffused but general activation at a time when controls display two well-ordered negative and positive domains, with a dynamic transition from the former to the latter. This was quantified by measuring virtual 4C contacts between either *Hoxd13*, *Hoxd11* or *Hoxd9* and the *Hoxd3-Hoxd4* region, which were markedly increased in mutant specimen (Fig. S12B).

In particular, when looking at contacts between the cluster and region 38-40 of sub-TAD1 (Fig. S13A, B), the step-wise extension of the loops towards more posteriorly located CBS observed in control (Fig. 3B, C) did not occur in mutant specimen. Instead, contacts were scored throughout the cluster, already from an early stage onwards, which translated into a bar rather than a spot in the CHi-C map (Fig. S13A, B). Altogether, these results indicated that CTCF proteins are both mandatory to protect posterior *Hoxd* genes to be contacted prematurely (Fig. S13C) and essential in organizing the precision and the pace of the *Hox* timer, even though remnant of a colinear process was observed in their almost complete absence.

## DISCUSSION

How developmental timing is both encoded and implemented remains a challenging question (Duboule, 2003; Ebisuya and Briscoe, 2018), as well as how evolutionary transitions may have been caused by relative modifications in temporal processes (see (Gould, 1977). During axial extension, the segmentation clock operates with a recurrent structure (Cooke and Zeeman, 1976; Palmeirim et al., 1997), driven by an oscillatory mechanism acting in-*trans* (Goldbeter and Pourquié, 2008). In contrast the *Hox* timer is a non-recurrent process, based on a *cis*-acting mechanism that uses the clustered organization of *Hox* genes to impose a time-sequence. Here, by using stembryos, we could document and analyze this process and propose a testable mechanistic model. We used stembryos (gastruloids (Turner et al., 2017) since they implement the *Hox* timer in a large proportion of their cells when compared to embryos, due to their prominent ‘posterior’ identities (Beccari et al., 2018; van den Brink et al., 2014; Veenvliet et al., 2020). Nonetheless the progressive restriction of expression domains towards the posterior aspect of the elongating stembryo (Beccari et al., 2018) dilutes any measured parameters and thus underestimates them, a bias for which we decided not to introduce any correction index. For example, *Hoxd9* is expressed in ca. 50 percent of cells expressing *Hoxd4* and thus any specific ‘positive’ signals (Pol II, H3K27ac, interaction bin) for *Hoxd9* is artificially reduced when compared to *Hoxd4*. This is particularly important to consider when small increases are observed, such as for example the progression of Poll II Ser2-p over *Hoxd9* at 144h, which are thus under-estimated in this study.

### The *Hox* timer; A three-steps mechanism

We propose a three steps mechanism for the *Hox* timer. A first step of *condensation* where the *Hox* clusters are packed into a negative globule, covered with H3K27me3 marks. While this had been observed previously in ES cells (Noordermeer et al., 2011; Soshnikova and Duboule, 2009), stembryos revealed that a central part of the cluster containing CBS1 loops out of this negative globule and is tightly anchored to several CBS located within the flanking sub-TAD1, prefiguring the two micro-domains to come, as if the cluster would be already in a pre-activation configuration. During the second *translocation* step, a quasi-simultaneous activation of the CTCF-free region is paralleled by a poorly colinear distribution of Pol II, which was an argument raised against the existence of a *Hox* timer by studying amphibians (Kondo et al., 2019). However, ChIP-seq of the elongating form of Pol II revealed a colinear distribution. From this, we conclude that a paused form of Pol II is deposited rapidly over the gene cluster (up to *Hoxd10*) and that its Ser2 phosphorylation occurs in a time-sequence.

During this *translocation* phase, the anterior part (up to CBS1) rapidly dissociated from the posterior part and cohesin complexes were loaded right onto this early transcribed part (Busslinger et al., 2017; Kagey et al., 2010; Zuin et al., 2014), with loop-extrusion mostly blocked by CBS1, as deduced from its preferential accumulation of RAD21. Soon after, extrusion involved more posteriorly located CBS, a progressive spreading clearly documented by our time-course interaction profiles, thus entering the third *progression* phase, also suggested by the change in the relative distribution of RAD21 over the CTCF sites. Interactions between more posterior genes and sub-TAD1 increased, likely as a result of extrusion reaching more posterior CTCF sites and bringing these genes in contact with potential sub-TAD1 enhancers.

Whether the mechanism of progression, i.e., what makes loop extrusion going through CBS1 at some point to reach CBS2, is active or passive is hard to evaluate. A passive process where extrusion would go through CBS1 either due a punctual lack of CTCF binding or to a leakage in the blocking efficiency of a bound CTCF, in both cases at a given frequency, could certainly account for the progression (Gabriele et al., 2022). Indeed, each event of this kind leading to the transcription of the gene located more ‘posteriorly’ would recruit cohesin at the promoter of the newly transcribed gene (Busslinger et al., 2017; Zhu et al., 2021) and thus promote extrusion from a more posterior position. After a certain time, cohesin loading would have progressed towards more posterior genes, along with their transcription, while still being loaded in 3’ of CBS1. Alternatively, the progression could be actively regulated by some CTCF cofactors such as MAZ, which was shown to contribute to CTCF insulation in the *HoxA* cluster (Ortabozkoyun et al., 2022) or by the WAPL binding partner PDS5 that also contributes to boundary function (Wutz et al., 2017).

The analyses of epigenetic profiles indicate that the other *Hox* gene clusters implement the exact same type of regulation. In this context, the extraordinary conservation in the number, positions and orientations of CTCF sites amongst paralogous mammalian clusters (i.e., over several hundred million years) (Amândio et al., 2021), or even when compared to birds (Yakushiji-Kaminatsui et al., 2018), can find an explanation. In particular the essential positioning of CBS1, which is found at the same relative position in all four *Hox* clusters, as well as the rather regular spacing between these CTCF sites, a feature that had remained unexplained when considered only in the context of a TAD boundary, which does not require such an iterated organization (e.g. (Anania et al., 2021).

### Heterochronic mutations and constrained flexibility

Our results also suggest that homeotic transformations previously observed in mice lacking CTCF sites in *Hox* clusters (Amândio et al., 2021; Narendra et al., 2016) may result from heterochronies, i.e., a wrong or unprecise timing in the activation of this series of genes. Thus far, however, no mutation or deletion was able to totally abolish the *Hox* timer such that *Hoxd13* would be transcribed concomitantly with *Hoxd1*, a situation lethal for the animal (Young et al., 2009; Zakany et al., 2004). We believe this is due to the initial transcriptional asymmetry, whereby *Wnt* signaling impacts *Hox* genes located in the ‘anterior’ part of the cluster (Neijts et al., 2016). From this and the subsequent enrichment in cohesin deposition in this region, more centrally and posteriorly located genes will become activated with some time delay, after they would exit the negative domain along with extrusion. In this view, the series of CTCF sites would fix the precision of the process as well as its proper sequence and pace, rather than being strictly mandatory for a global time delay in transcription to occur.

From an evolutionary viewpoint, the separation between an intrinsic tendency to produce a time sequence in the activation of series of genes in-*cis* due to an asymmetric transcriptional activation and the mechanism regulating the pace and the precision of this mechanism makes sense, for it allows to customize the outcome in a species-specific manner. Indeed, a slight variation in the timing of activation may have important consequences upon axial structures and such variations might have been essential to produce the distinct *Hox* combinations observed in different vertebrates (Burke et al., 1995; Gaunt, 1994). For instance, changes in the number of CTCF sites, their positions or even their binding affinities may have triggered modifications in the expression timing of *Hox* genes. However, the presence of four *Hox* gene clusters containing similarly organized CTCF sites also makes the overall collinear process robust and resilient to potential perturbations. This buffering effect and hence the evolutionary stability of the homeotic system within any given species, is likely re-enforced by the subsequent use of subsets of these CTCF sites for different purposes, after the *Hox* timer is over, for example for the determination of cell types (Narendra et al., 2015) or to position other enhancers located in the flanking TADs, which are necessary for the many functions *Hox* genes fulfill after their axial contribution is achieved (Amândio et al., 2021; Rodríguez-Carballo et al., 2020).

## MATERIALS AND METHODS

### Culture of stembryos/gastruloids

Mouse embryonic stem (mES) cells were routinely cultured in gelatinized tissue-culture dishes with 2i (Silva et al., 2008) LIF DMEM medium composed of DMEM + GlutaMAX supplemented with 10% ES certified FBS, non-essential amino acids, sodium pyruvate, beta-mercaptoethanol, penicillin/streptomycin, 100 ng/ml of mouse LIF, 3 µM of GSK-3 inhibitor (CHIR99021) and 1 µM of MEK1/2 inhibitor (PD0325901). Cells were passaged every 3 days and maintained in a humidified incubator (5% CO2, 37 °C). The differentiation protocol for gastruloids was previously described (Beccari et al., 2018). Briefly, ES cells were collected after accutase treatment, washed and resuspended in prewarmed N2B27 medium (50% DMEM/F12 and 50% Neurobasal supplemented with 0.5x N2 and 0.5x B27). 300 cells were seeded in 40 µl of N2B27 medium in each well of a low-attachment, rounded-bottom 96-well plate. 48h after aggregation, 150 µl of N2B27 medium supplemented with 3 µM of GSK-3 inhibitor was added to each well. 150 µl of medium was then replaced every 24h. Collection of gastruloids for each time-point was performed indiscriminately up to 96h after aggregation. Starting from 108h, only those gastruloids showing a clear elongating shape with no sign of apoptosis were collected and processed for subsequent analysis.

Because gastruloids grown under various protocols in various laboratories can be very different from one another (e.g. (Beccari et al., 2018; van den Brink et al., 2020; Veenvliet et al., 2020), we refer to them as stembryos throughout the paper to prevent confusions.

### Generation of mutant mES cells

Wild type mES cells (EmbryoMax® 129/SVEV) were used to generate mutant cell lines following the CRISPR/Cas9 genome editing protocol described in (Andrey et al., 2017; DOI 10.1007/978-1-4939-4035-6_15). sgRNA targeting guides (Supplementary Table S1) were cloned into a Cas9-T2A-Puromycin expressing plasmid containing the U6-gRNA scaffold (gift of Andrea Németh, Addgene plasmid #101039). mES cells were transfected with 8 µg of plasmid using the Promega FuGENE 6 transfection kit and dissociated 48h later for puromycin selection (1.5 µg/ml). Clone picking was conducted 5 to 6 days later and positive mES cell clones were assessed by PCR-screen using the MyTaq PCR mix kit (Meridian bioscience) and specific primers surrounding the targeted region (Supplementary Table S2). Mutations were verified for both alleles by sanger sequencing (Supplementary Table S3). When heterozygous mutations were obtained, the transfection procedure was reiterated until homozygous mES cell clones were obtained. For the Del(CBS1-2) mutant, the deletion of CBS2 was carried out on top of Del(CBS1) ES cells. The Ins(2xCBS-d4d8) cell line was produced by the insertion of a 900 bp DNA cassette at 2 kb distance 5’ to CBS1. The cassette contains 2 CTCF sites with the same orientation as CBS1. The transfection was performed with 1.5 µg of the plasmid containing the recombination cassette along with 8 µg of the sgRNA plasmid. The Del(CBS1-5) and Del(sub-TAD1) ES cell lines were derived from mouse blastocysts at the Mouse Clinical Institute (www.ics-mci.fr). The Del(CBS1-5) mouse was described in (Amândio et al., 2021). Del(sub-TAD1) mice were obtained by electroporating the CRISPR guide (Supplementary Table S1) and the Cas9 mRNA into fertilized mouse embryos. The deletion covers the sub-TAD1 region including *Mtx2* gene, but not the CS38-40 boundary. To overcome *Mtx2* knock out homozygous lethality, mice carrying as a transgene a modified murine fosmid containing a sequence covering the *Mtx2* region (based on RP24-284D11 from https://bacpacresources.org, see (Allais-Bonnet et al., 2013)) were crossed with the Del(sub-TAD1) mice for rescue.

### NGS analysis and figure generation

All NGS analyses were performed on a local installation of galaxy (Afgan et al., 2016). All command lines and scripts to regenerate figures are available on https://github.com/lldelisle/scriptsForRekaikEtAl2022. Most of genomic tracks were plotted using pyGenomeTracks version 3.7 (Lopez-Delisle et al., 2021; Ramirez et al., 2016) and modified with illustrator. All boxplots and barplots were plotted using Prism, heatmaps were plotted with R (www.r-project.org).

### Mutant genomes

Chromosome 2 matching the 3 mutants (Ins(2xCBS-d4d8), Del(d1-d4) and Del(sub-TAD1) were generated manually using the chromosome 2 sequence from UCSC of mm10 and results from sanger sequencing of the breakpoints (Rekaik and Lopez-Delisle, 2022). The mutant chromosome 2 sequence was concatenated with the sequences of all other autosomes, chrX, chrY and mitochondrial DNA from mm10 (UCSC).

### RNA-seq

Stembryos for each condition were collected and pooled in a 2 ml Eppendorf tube. After centrifugation and medium removal, pelleted stembryos were stored at −80°C until RNA extraction. RNeasy Plus Micro kit (Qiagen) with on-column DNase digestion was used for RNA extraction following the manufacturer’s instructions. RNA-seq library preparation with Poly-A selection was performed with 1 µg of purified RNA using the TruSeq Stranded mRNA kit from Illumina and following the manufacturer’s protocol. Library quality was assessed with Fragment analyzer before sequencing on a NextSeq 500 sequencer as paired-end, 75 bp reads. For data analysis, adapter sequences were trimmed from reads using cutadapt (Martin, 2011) version 1.16 (-a ’GATCGGAAGAGCACACGTCTGAACTCCAGTCAC’ -A ’GATCGGAAGAGCGTCGTGTAGGGAAAGAGTGTAGATCTCGGTGGTCGCCGTATCATT’). Trimmed reads were aligned on mm10 with STAR version 2.7.7a (Dobin et al., 2013) with ENCODE options using a custom gtf based on Ensembl version 102 (Lopez-Delisle, Lucille, 2021). Only uniquely mapped reads (tag NH:i:1) were kept using bamFilter version 2.4.1 (Barnett et al., 2011). FPKM values were computed using cufflinks version 2.2.1 (Roberts et al., 2011; Trapnell et al., 2010) with default parameters. In order to reduce the impact of variations in gastruloid growth speed, the FPKM values of *HoxD* genes (except *Hoxd13*) and some *HoxA* genes (*Hoxa1*, *Hoxa5* and *Hoxa9*) were normalized using FPKM values of genes from other clusters (*Hoxa* and *Hoxc* genes were used to normalize *Hoxd* whereas *Hoxb* and *Hoxc* genes were used to normalize *Hoxa*) (see github repository for more details).

### ChIP and ChIPmentation (ChIP-M)

ChIP and ChIP-M experiments were performed according to the protocol described in (Darbellay et al., 2019; Rodriguez-Carballo et al., 2017) and adapted for stembryos samples. Briefly, collected stembryos were pooled in a 15ml falcon tube, washed with PBS and resuspended in 1ml PBS containing 1% formaldehyde for fixation during 10 min at room temperature. The crosslink reaction was stopped by adding a glycine solution to a final concentration of 0.125 M. Fixed stembryos were pelleted and stored at −80°C until further use. Samples were resuspended in a sonication buffer (Tris HCl pH=8.0 50 mM; EDTA 10 mM; SDS 0.25% and protease inhibitors) and sonicated in a Covaris E220 device for 14 min (duty cycle 2%, peak incident power 105 W) to obtain an average chromatin fragment size of 300-500 bp. A dilution buffer (HEPES pH=7.3 20 mM; EDTA 1 mM; NP40 0.1%; NaCl 150 mM and protease inhibitors) was added to the sonicated chromatin and incubated with the antibody-bead complex (Pierce Protein A/G Magnetic Beads, Thermo Scientific) overnight at 4°C. Sequential washes were then performed twice with RIPA buffer (Tris HCl pH=8.0 10 mM; EDTA 1 mM; Sodium Deoxycholate 0.1% TritonX-100 1%; NaCl 140 mM and protease inhibitors), RIPA High salt buffer (Tris HCl pH=8.0 10 mM; EDTA 1 mM; Sodium Deoxycholate 0.1% TritonX-100 1%; NaCl 500 mM and protease inhibitors), LiCl buffer (Tris HCl pH=8.0 10 mM; EDTA 1 mM; LiCl 250 mM; Sodium Deoxycholate 0.5%; NP40 0.5% and protease inhibitors) and Tris HCl buffer (pH=8.0 10 mM and protease inhibitors). For ChIP experiments [H3K27ac (ab4729, Abcam), H3K27me3 (39155, Active Motif) and PolII-Ser2p (04-1571, Millipore)], DNA fragments were incubated in the elution buffer (Tris HCl pH=8.0 10 mM; EDTA 5 mM; NaCl 300 mM and SDS 0.1%) containing proteinase K and purified using the Qiagen MiniElute kit. A phosphatase inhibitor (PhosStop, Roche) was used during PolII-Ser2p ChIP incubation and beads wash. The DNA library was produced using TruSeq adapters and amplified with the Kapa HiFi library kit (Roche) using a number of cycles determined by qPCR. For ChIP-M samples [CTCF (61311, Active Motif), PolII (ab817, Abcam), RAD21 (ab992, Abcam) and NIPBL (A301-779A, Bethyl Laboratories)], DNA fragments bound to the antibody-bead complex were tegmented using the Nextera Tegmentation kit. Beads were resuspended in the tegmentation buffer and incubated at 37° for 2 min with 1 µl of the Tn5 transposase. Fragment were then eluted and purified as described previously, and amplified using Nextera primers. Final DNA libraries were purified and size selected using AMPure XP magnetic beads (Beckman Coulter), and a fragment analysis was performed before sequencing on a NextSeq 500 sequencer as paired-end, 75 bp reads. For data analysis, fastqs of the 3 replicates of wt_144h_pSer2PolII were concatenated before any processing. Adapter sequences and bad quality bases were trimmed from reads using cutadapt (Martin, 2011) version 1.16 (-a ’GATCGGAAGAGCACACGTCTGAACTCCAGTCAC’ -A ’GATCGGAAGAGCGTCGTGTAGGGAAAGAGTGTAGATCTCGGTGGTCGCCGTATCATT’ for ChIP and -a ’CTGTCTCTTATACACATCTCCGAGCCCACGAGAC’ -A’CTGTCTCTTATACACATCTGACGCTGCCGACGA’ for ChIPmentation --quality-cutoff=30). Trimmed reads were mapped on mm10 genome (or the mutant genome Ins(2xCBS-d4d8) for the corresponding mutant CTCF ChIPmentation) using bowtie2 version 2.3.4.1 (Langmead and Salzberg, 2012). Only alignments with proper pairs and mapping quality above 30 were kept using samtools 1.8. Peaks and coverage were obtained with macs2 version 2.1.1.20160309 (--format BAMPE --gsize 1870000000 --call-summits --bdg). In order to better compare ChIP tracks, ChIP samples for the same protein with a time-course were normalized together using a custom python script (available in the github repository). Similarly, this normalization was used when mutant and control were performed in parallel (see github repository). ChIP quantifications were performed with multiBigWigSummary from deeptools version 3.0.0 (Ramirez et al., 2016). The cumulative H3K27ac profile of figure 2A was computed with a custom python script available on the github repository. In order to reduce the impact of variations in gastruloid growth speed for the H3K27ac and H3K27me3 datasets in supplementary figure S9A and supplementary figure S12A, the profiles of mutant and corresponding wild-type were corrected using the time-course experiment and the profile at the *HoxA* cluster as a guide (see github repository for more details). The profiles were then averaged for the two replicates and differences were performed using bash script available in github repository.

### Capture Hi-C

Collected stembryos were pooled and fixed for 10 min in 1% formaldehyde at room temperature and stored at −80°C until further use. Lysis was performed with TissueLyser II and metallic beads for 30 sec at 3hertz before incubation at 4° for 30 min. The rest of the protocol for the capture Hi-C library generation is described in (Bolt et al., 2021). The customized SureSelectXT RNA probe used for DNA capture cover the region from chr2:72240000 to 76840000 (mm9). For data analysis, TruSeq adapters and bad quality bases were removed with cutadapt (Martin, 2011) version 1.16 (-a ’AGATCGGAAGAGCACACGTCTGAACTCCAGTCAC’ -A ’AGATCGGAAGAGCGTCGTGTAGGGAAAGAGTGTAGATCTCGGTGGTCGCCGTATCATT’ --minimum-length=15 --quality-cutoff=30). Trimmed reads were processed with HiCUP version 0.6.1 (Wingett et al., 2015) with default paramters on mm10 or on the corresponding mutant genome. The output BAM was converted to Juicebox format thanks to a custom python script available in (https://testtoolshed.g2.bx.psu.edu/repository/download?repository_id=be5040251cd4afb7&changeset_revision=44365a4feb3b&file_type=gz). Only pairs with both mates falling into the capture region (mm10:chr2:72402000-77000000) and with both mates with quality above 30 were kept. Filtered pairs were loaded into 2kb and 5kb matrices with cooler version 0.7.4 (Abdennur and Mirny, 2020). The matrices were balanced with cooler balance (--cis-only). For the Del(sub-TAD1) mutant, the balancing was done with the option --ignore-diags 5 instead of --ignore-diags 2 to discard the bias due to the presence of a BAC containing the region (mm10:chr2:74747769-74913171). On figure 3C, the histograms were produced with Plot profile on imageJ version 2.1.0/1.53c. Virtual 4C were computed using the same script as in (Amândio et al., 2021) inspired by (Despang et al., 2019). The output bedgraph were then used for the quantifications.

### HiChIP

Wild type stembryos were collected at 120h and fixed for 10 min in 1% formaldehyde at room temperature before proceeding with the library generation. The protocol for H3K27ac HiChIP was adapted from (Mumbach et al., 2016). Briefly, after lysis, DpnII digestion, biotin fill-in and blunt-end ligation (see Capture Hi-C), the samples were sonicated in a Covaris E220 instrument for 10 min (duty cycle 2%, peak incident power 105 W) and incubated overnight at 4° with magnetic beads conjugated to the H3K27ac antibody (Abcam, ab4729). The next day, beads were washed as described in ChIP protocol before elution of DNA fragments. A biotin pull-down was then performed (Dynabeads MyOne Streptavidin T1) followed by a TruSeq library preparation using fragments bound to the streptavidin beads. A qPCR was used to estimate library depth before final amplification with the Kapa HiFi library kit. Library was purified, size selected and quality assessed with fragment analyzer before sequencing on a NextSeq 500 sequencer as paired-end, 75 bp reads. Hi-ChIP reads were processed similarly to capture Hi-C, except that the adapters were removed on each read independently and bad quality pairs were conserved, pairs were not filtered for the capture region, pairs were loaded into 10 kb bins and no balancing was performed.

### Generation of supplementary movies

For movie 1, the H3K27ac profiles of figure 1C were quantified on each bin of 10 bp. Between every measured timepoints (12h time-course) intermediate timepoints (every hour) were generated by linear extrapolation on each bin using the previous and the next time points such as to smoothen the profiles and to see their dynamic. Profiles were then generated by pyGenomeTracks and combined in a film with ImageJ.

For movie 2, the balanced, 5 kb-bins, cool files used in figure 3A, B were used to generate intermediate cool files for each hour using a linear extrapolation on each 5 kb bin using the previous and the next time point. Profiles were then generated by pyGenomeTracks. Annotations (arrows, bars) were manually added and all frames were combined in a film using ImageJ.

## ACKNOWLEDGMENTS

We thank Dr. Fabrice Darbellay for advice with ChIPmentation experiments, Chase Bolt for his assistance with Capture Hi-C experiments, Marie Anselmet for her help with RNA-seq experiments as well as other colleagues from the Duboule laboratories for discussions. We are grateful to Anne-Catherine Cossy for her technical assistance with cell culture. This work was supported in part using the resources and services of the Gene Expression Research Core Facility (GECF) at the School of Life Sciences of EPFL.

## AUTHORS CONTRIBUTIONS

Conceptualization: HR, DD

Methodology: HR, LLD, AH, BM, CB

Investigation: HR, CB, AH

Visualization: HR, LLD

Funding acquisition: DD

Project administration and supervision: DD

Writing the original draft: HR, DD

Review writing & editing: HR, DD, LLD

## DATA AND MATERIALS AVAILABILITY

All raw and processed datasets are available in the Gene Expression Omnibus (GEO) repository under accession number GSE205783. All scripts necessary to reproduce figures from raw data are available at https://github.com/lldelisle/scriptsForRekaikEtAl2022

## SUPPLEMENTARY MATERIALS

### SUPPLEMENTARY FIGURES S1 TO S13 AND LEGENDS

**Supplementary Figure 1.**
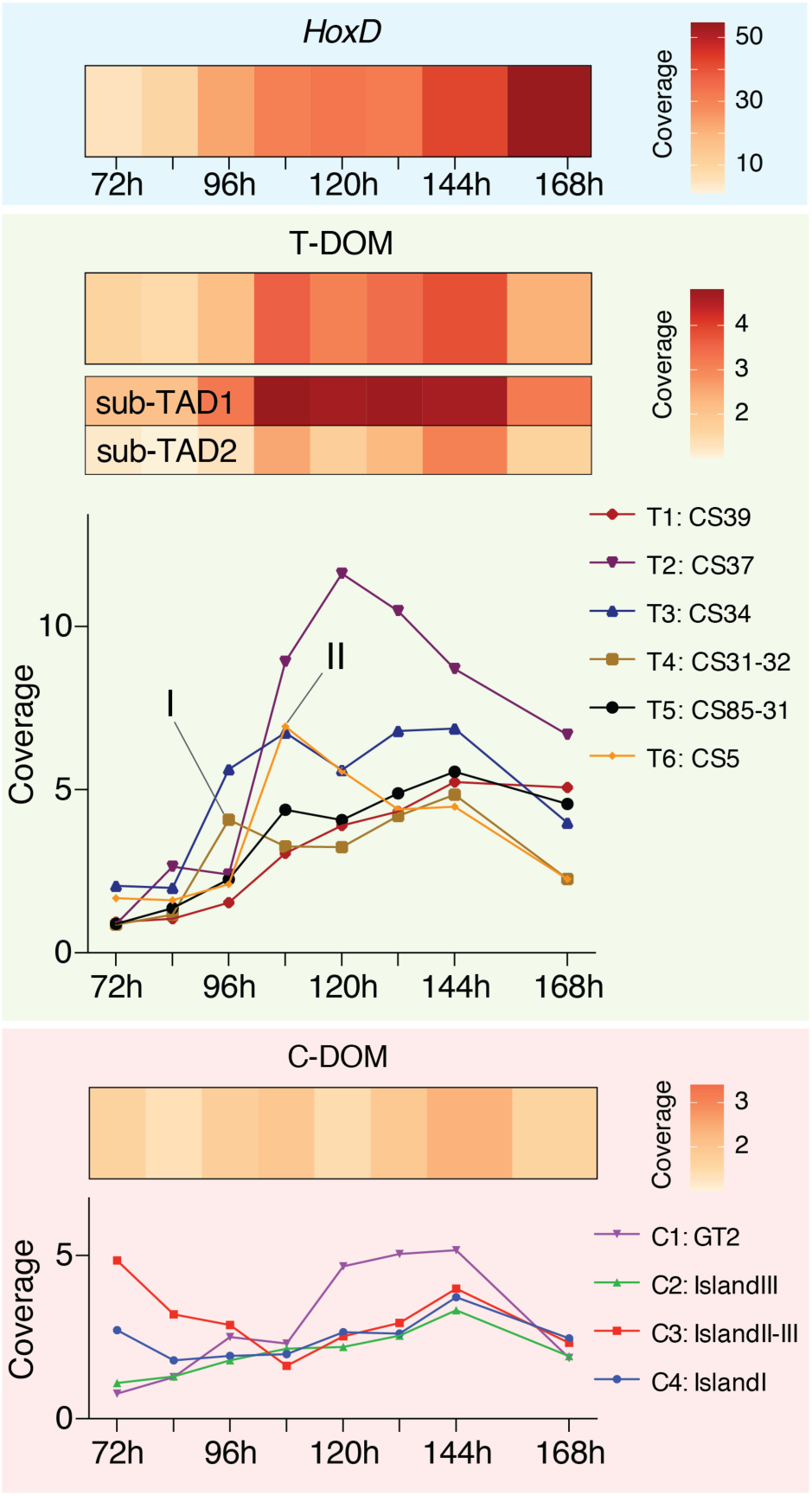
Quantification of the H3K27ac ChIP-seq coverage over the *HoxD* cluster (top), the T-DOM TAD (middle) and the C-DOM TAD (bottom). Each square represents the average coverage value for a given time-point. The increase in H3K27ac coverage over the *HoxD* cluster and T-DOM (with a strong preference for sub-TAD1) contrasts with C-DOM (bottom). Graphs showing quantifications over time of the colored vertical segments illustrated in Fig. 1B (red for C-DOM and green for T-DOM). Each line represents a specific H3K27acetylated element within the *HoxD* regulatory landscape. The time sequence in the activation of these elements in T-DOM does not correlate with their distance from the genes cluster (I and II).

**Supplementary Figure 2.**
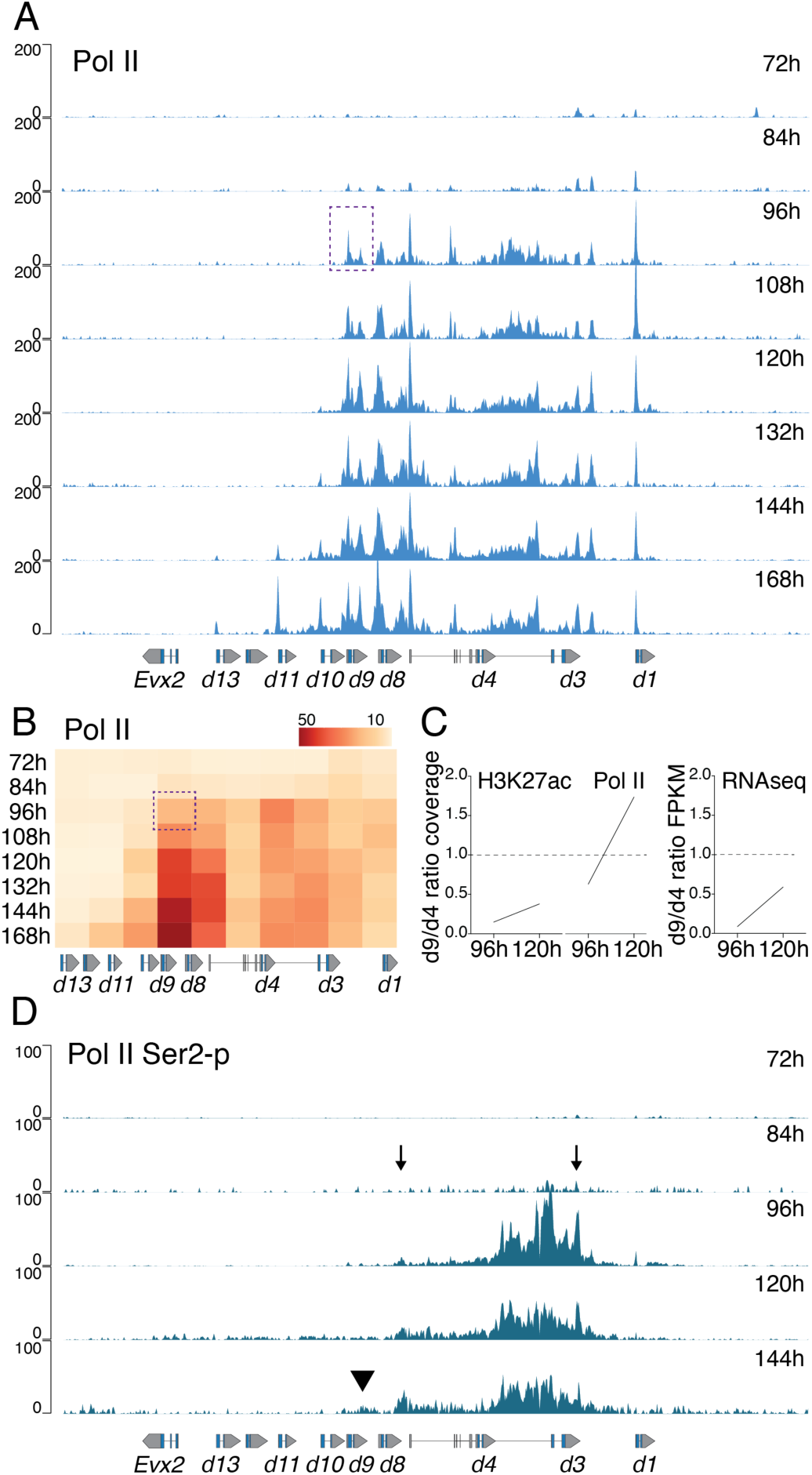
Pol II distribution during the activation of the *HoxD* cluster. (A) Pol II ChIP-seq profiles over *HoxD* at different stembryonic stages. While signals are enriched over *Hoxd1* and *Hoxd3* at 72h and 84h, a surprisingly fast and large coverage in Pol II is scored at 96h, up to *Hoxd9* (dashed region). (B) Heatmap of Pol II ChIP-seq coverage over the *HoxD* cluster divided into 10 kb bins. (C) Computed ratio between *Hoxd9* (dashed square) and *Hoxd4* coverage (fourth bin from *Hoxd1*) for both H3K27ac and Pol II at 96h and 120h (left panel), and between their FPKM values (right panel). Comparable ratio between H3K27 acetylation and transcription was observed, unlike with Pol II where enrichment was already observed over *Hoxd9* at 96h (D) ChIP-seq profiles of Pol II phosphorylated at serine 2 at five time-points. The dynamic of the elongating Pol II Ser2-p was comparable to H3K27 acetylation though with a slight delay.

**Supplementary Figure 3.**
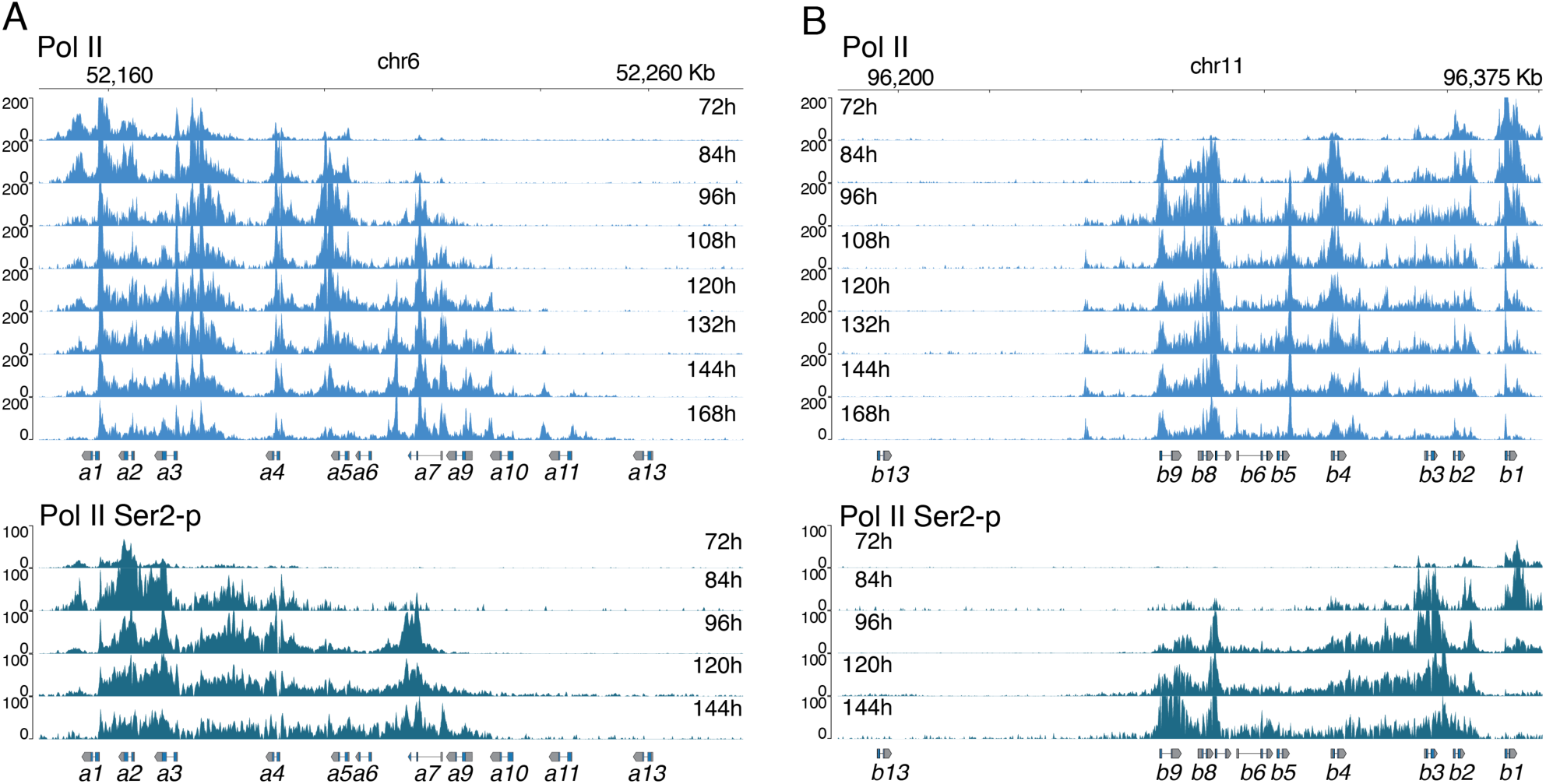
Pol II and Pol II Ser2P profiles on *HoxA* and *HoxB* clusters. On top a time course of Pol II ChIP-seq profiles over the *HoxA* (A) and *HoxB* (B) genes clusters in stembryos from 72h to 168h, showing a rapid spreading of Pol II over the gene clusters. Bottom: The use of an antibody against Pol II with phosphorylated serine 2 in ChIP-seq profiles at various time points reveals distinct, more progressive and colinear profiles, suggesting that Pol II is initially pausing over the gene clusters before starting to elongate.

**Supplementary Figure 4.**
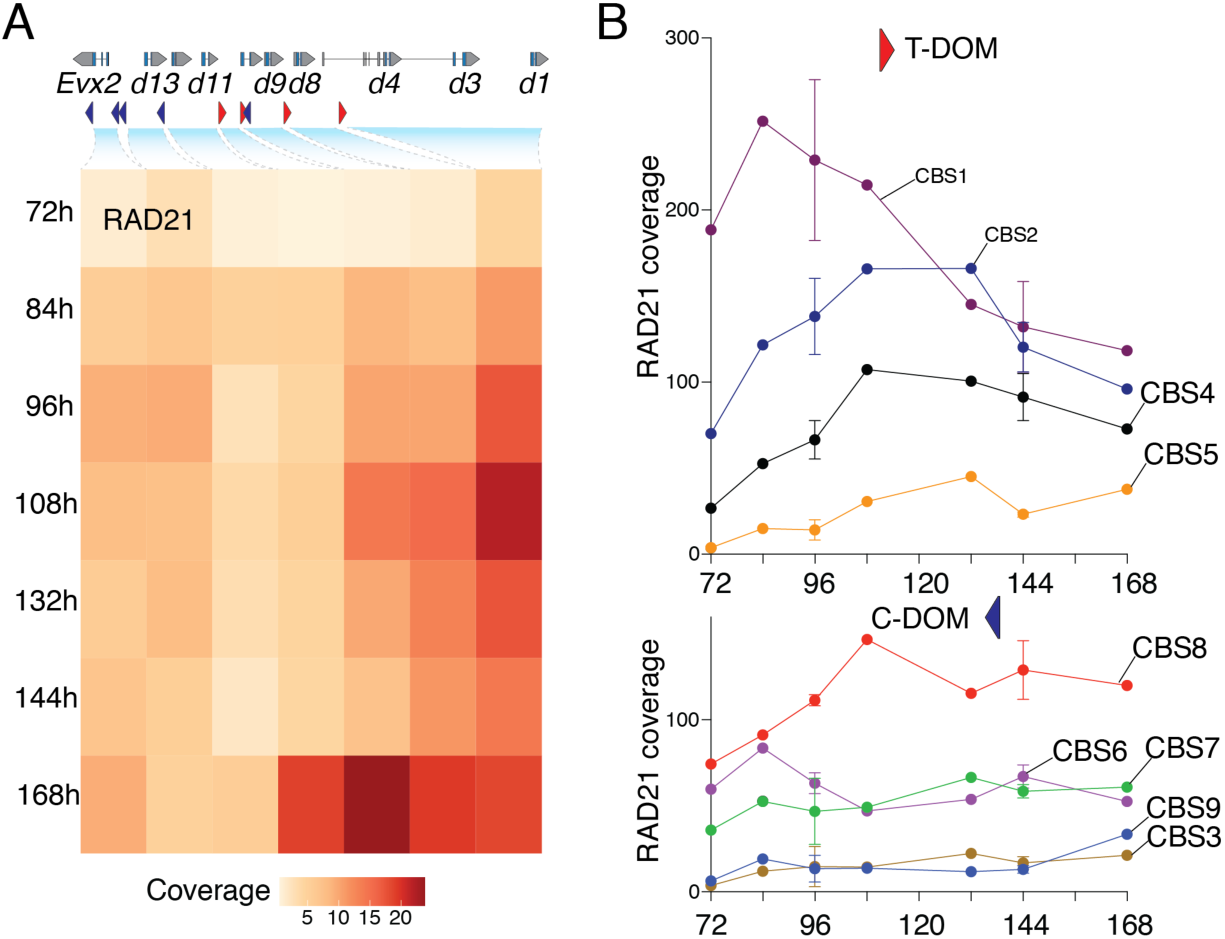
Quantification of RAD21 coverage. (A) Heatmaps of RAD21 ChIP-seq coverage over inter-CBS regions at the indicated time-points (n=1). Each square represents a segment located between two CBS within the *HoxD* cluster (scheme on top). The positions and orientations of CBS are indicated with red and blue arrowheads. An increased RAD21 coverage is detected over the activated region of the gene cluster, when compared to the silent posterior part. (B) RAD21 ChIP-seq coverage over those CBS located within the cluster and oriented either toward T-DOM (top), or towards C-DOM (bottom). The dynamic of coverage over time (x-axis) indicates a progressive accumulation of extruding cohesin at those CBS oriented toward T-DOM, i.e., where active loading was observed (n=1; 96 and 144h: n=2).

**Supplementary Figure 5.**
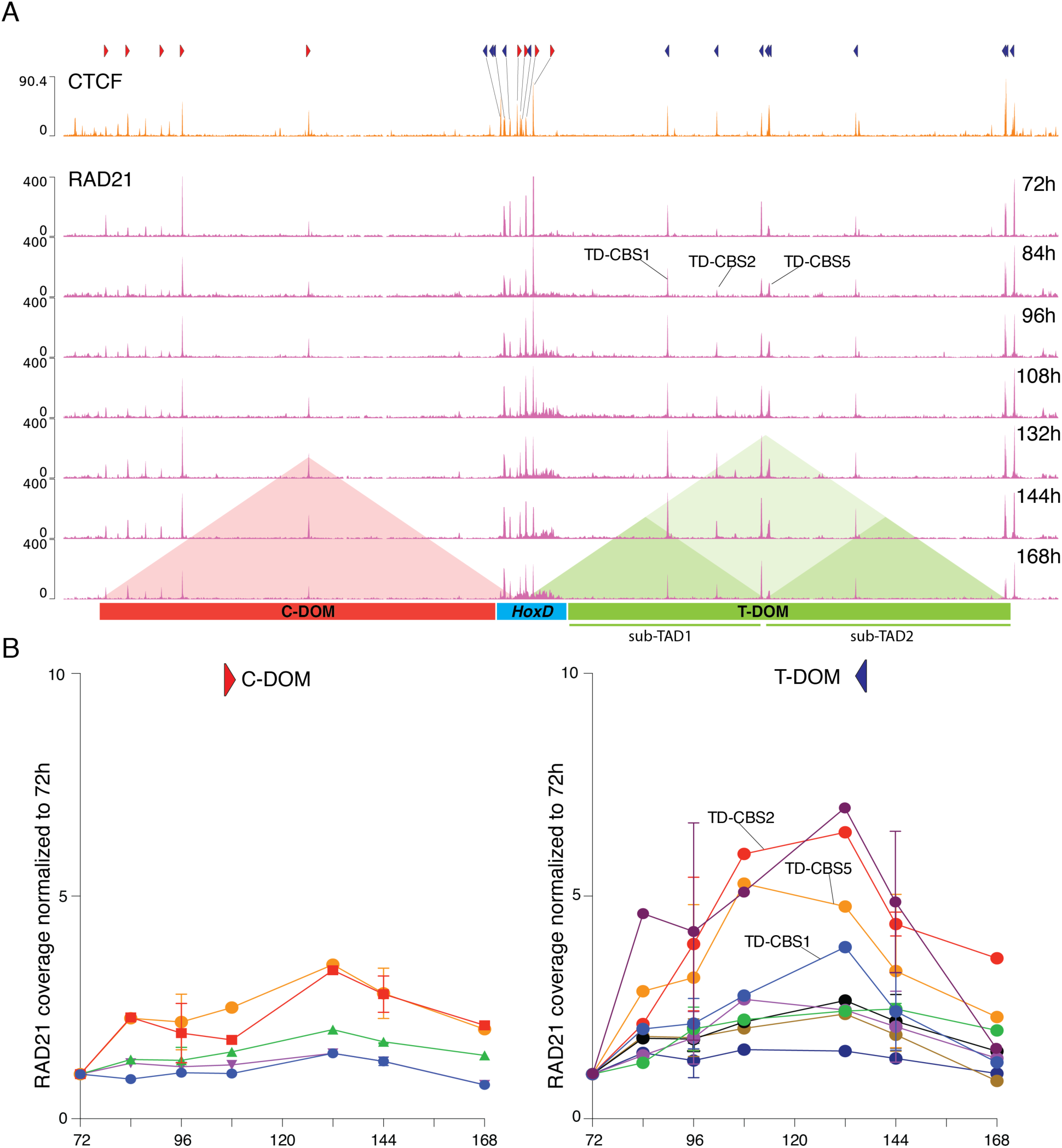
CTCF (A, top) and RAD21 (A, bottom) ChIP-seq profiles over the *HoxD* locus. The positions and orientations of CBS are indicated with red and blue arrowheads. The labeled CBS within the sub-TAD1 (on the 84h profile) are those involved with the translocation of the cluster. (B) RAD21 ChIP-seq coverage over those CBS located outside the cluster, either within T-DOM (right), or within C-DOM (left). The quantification of RAD21 coverage over time (x-axis) indicates a progressive accumulation of extruding cohesin at some CBS located within sub-TAD1 (n=1; 96 and 144h: n=2).

**Supplementary Figure 6.**
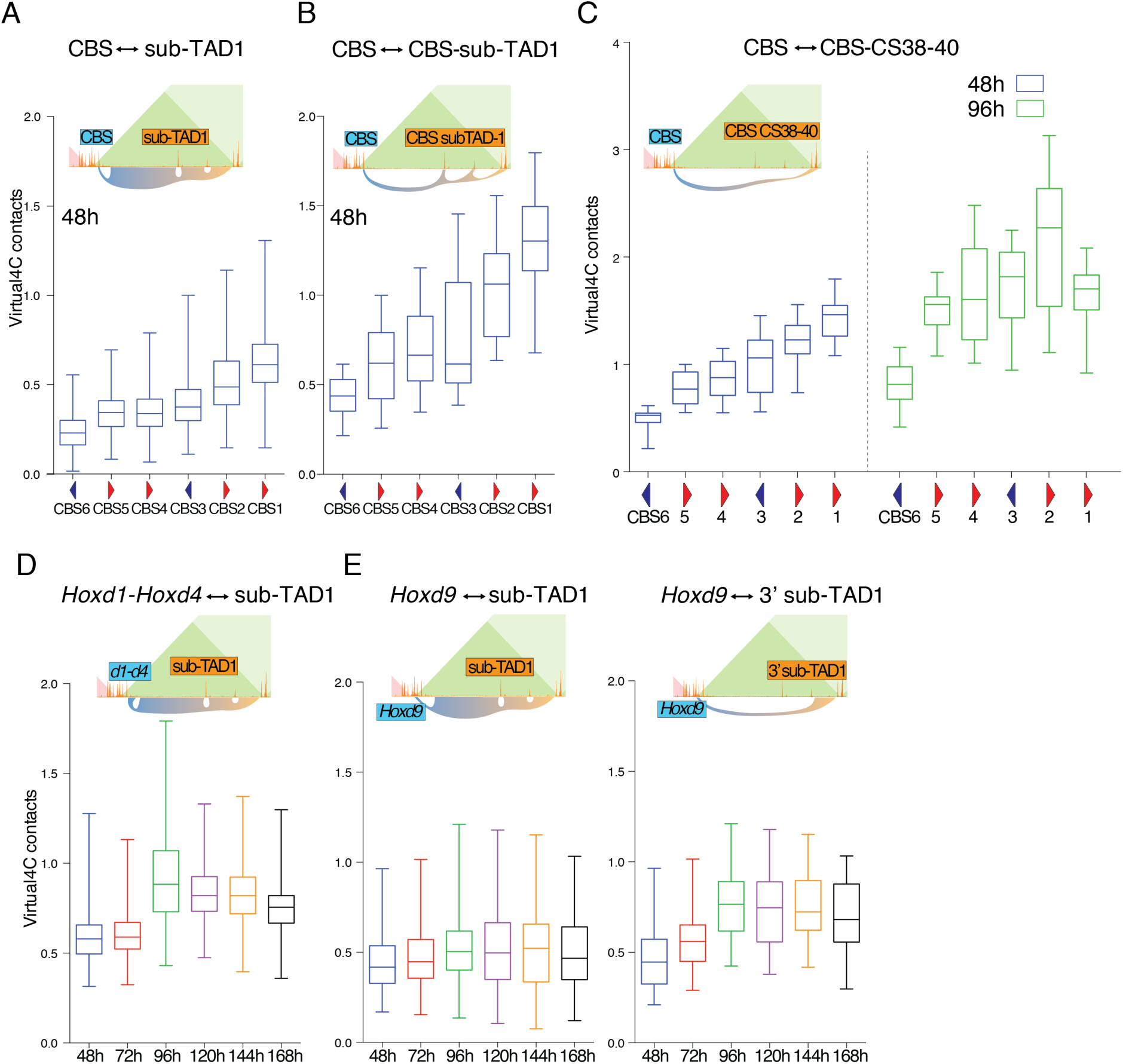
Virtual 4C contacts generated from Capture Hi-C datasets produced from control stembryos. For each quantification, a scheme depicts the viewpoint (blue) and the contact established (orange). (A) Quantification of contacts between the various CBSs in *HoxD* and regions of sub-TAD1 (sub-TAD1 CBSs excluded) at 48h. (B) Quantification of contacts between the various CBSs in *HoxD* and the CBSs located in sub-TAD1 at 48h. CBS1 establishes more contacts than any other CBS within *HoxD*, with sub-TAD1. (C) Quantification of contacts between the various CBSs in *HoxD* and those CBSs within the CS38-40 region at 48h and 96h, showing the preponderant contacts with CBS2 after activation (96h). (D) Contacts between the *Hoxd1-Hoxd4* region and sub-TAD1 increased upon cluster activation at 96h. (E) Contacts between *Hoxd9* and sub-TAD1 (left), or the 3’ region of sub-TAD1 only (right) at 96h show increased interactions with the 3’ part of the regulatory landscape upon activation of the cluster. Box plots with median value and 25-75% percentiles, whiskers represent minimum and maximum.

**Supplementary Figure 7.**
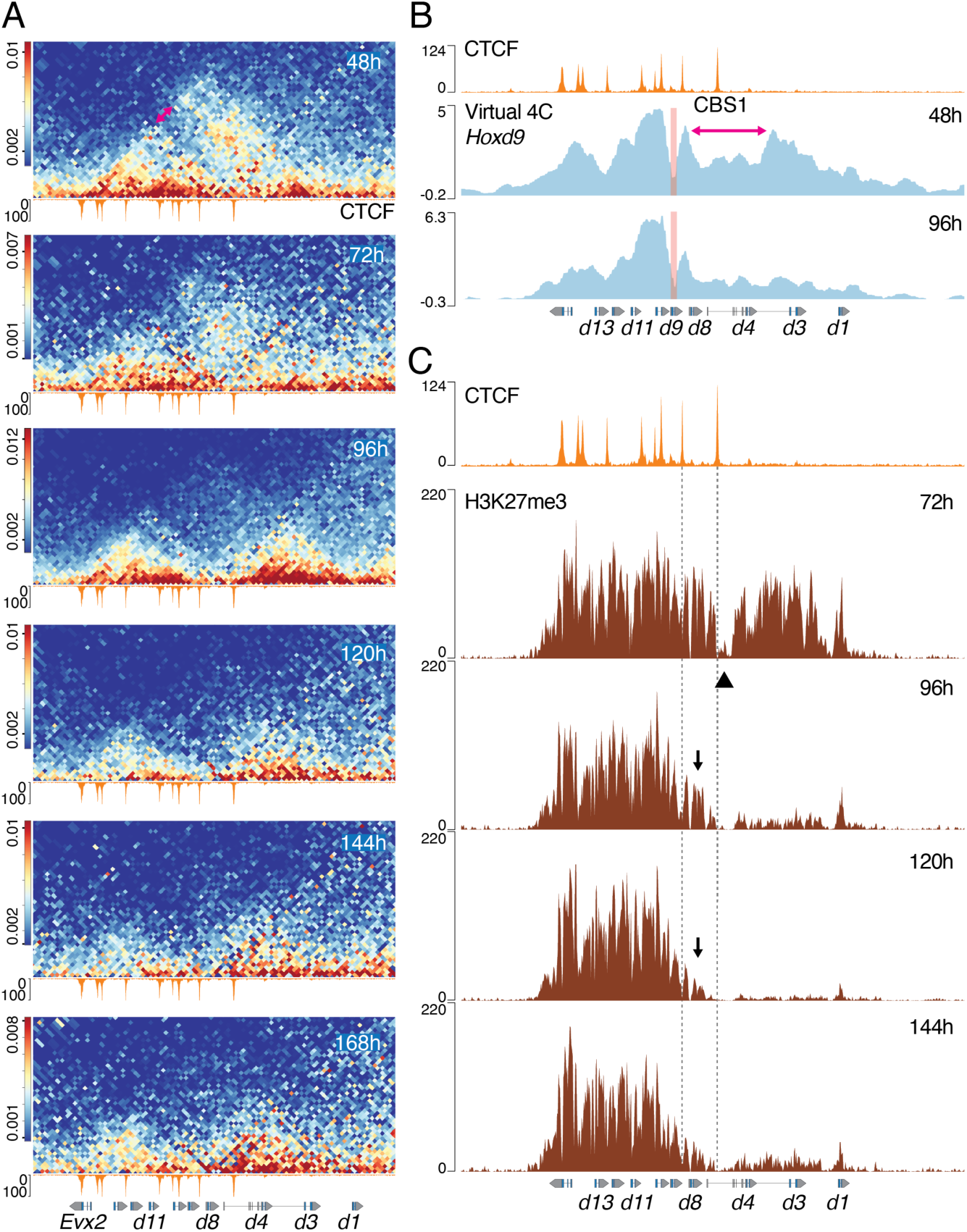
Dynamic reallocation of intra-cluster interactions. (A) Magnifications of the capture Hi-C map shown in Fig. 4, focusing on the interactions internal to *HoxD* at 2kb resolution. A CTCF ChIP track at 168h is shown below each panel and the genes are at the bottom. The double red arrow indicates the looping out of a central part of the cluster. Full translocation of the anterior part is observed at 96h and a weakening of the negative domain (left) is observed thereafter. (B) Virtual 4C contacts generated from Capture Hi-C data at 48h and 96h, using the *Hoxd9* gene body as a viewpoint (red vertical line). After translocation, the contacts with *Hoxd4*-*Hoxd1* are much reduced. The region of the cluster that initially loops out is indicated with a red arrow in (same as in panel A, top). (C) H3K27me3 ChIP-seq of stembryos at various time points, showing the activation of the anterior part, followed by a step-wise retraction (black arrows) up to 120h, when it appears stabilized (see the text). A CTCF ChIP-seq profile is on top for alignment. The fragment looping out is indicated with arrowhead. The two vertical lines help to position CBS1 and CBS2.

**Supplementary Figure 8.**
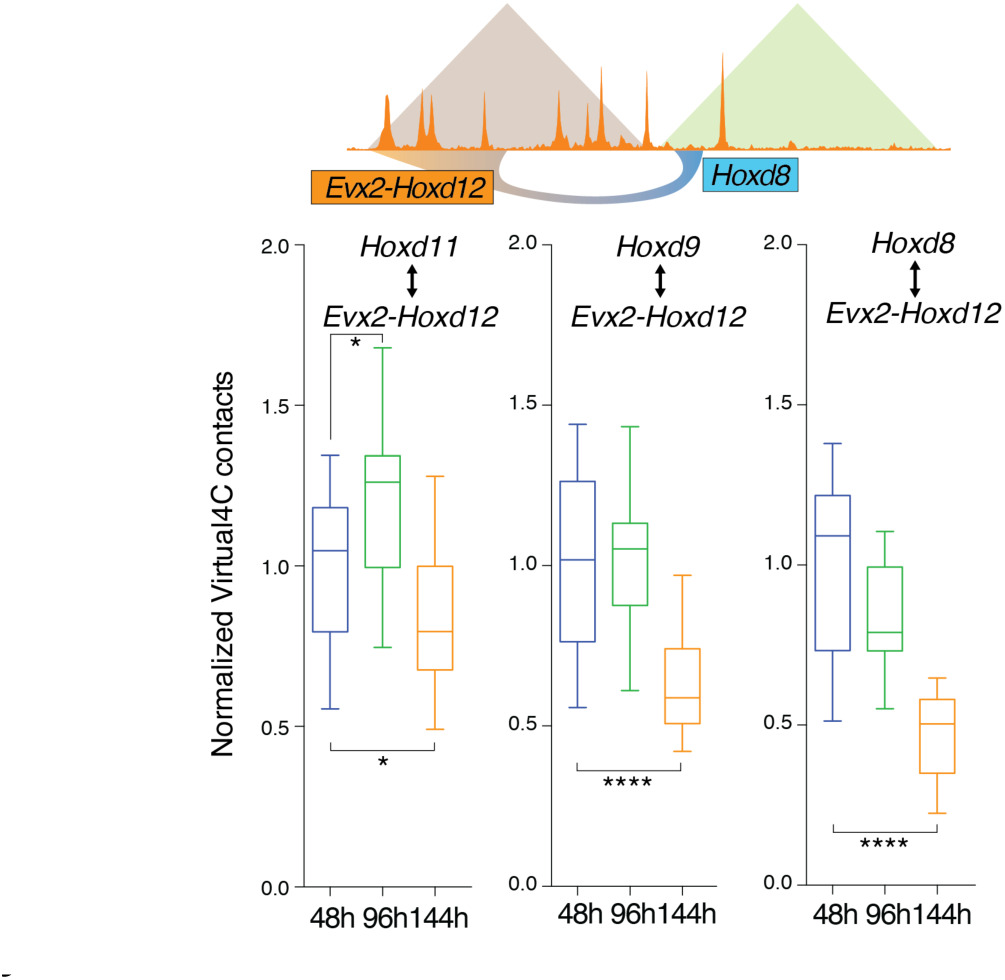
Virtual 4C contacts generated from Capture Hi-C datasets produced out of control stembryos at 48h, 96h and 144h. The contacts between the various viewpoints (*Hoxd* genes shown on top) and the *Evx2-Hoxd12* region showed a progressive decrease upon activation of the cluster, particularly visible for both *Hoxd8* and *Hoxd9*. Contacts values were normalized to 48h for each viewpoint. Asterisks represent significant changes against the 48h time-point as determined by the Kruskal-Wallis followed by Dunn’s multiple comparisons test (* is p-value < 0.05, **** < 0.0001). Box plots with median value and 25-75% percentiles, whiskers represent minimum and maximum.

**Supplementary Figure 9.**
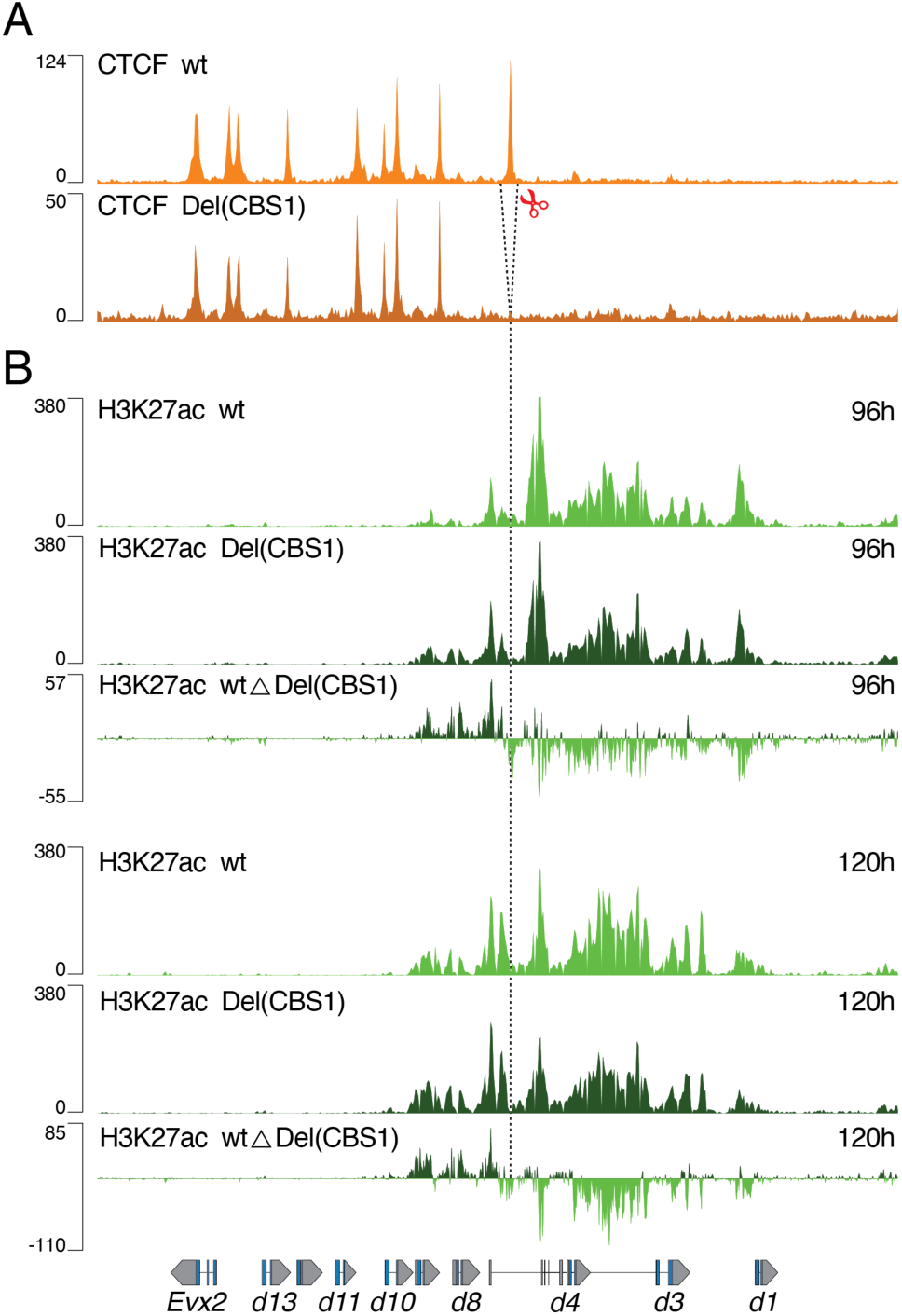
H3K27ac in stembryos lacking CBS1. (A) CTCF ChIP-seq of control stembryos at 168h (top) and Del(CBS1) stembryos at 96h (bottom). The deleted fragment containing CBS1 is indicated with scissors. (B) H3K27ac ChIP-seq using control and Del(CBS1) stembryos at 96h (top) and 120h (bottom)(average of two replicates). A subtraction track with acetylation coverage is shown for each time-point, with colors indicating either control (light green) or Del(CBS1) (dark green) enrichments. The vertical line indicates the position of deleted CBS1, i.e., exactly where the inversion of the enrichment occurred.

**Supplementary Figure 10.**
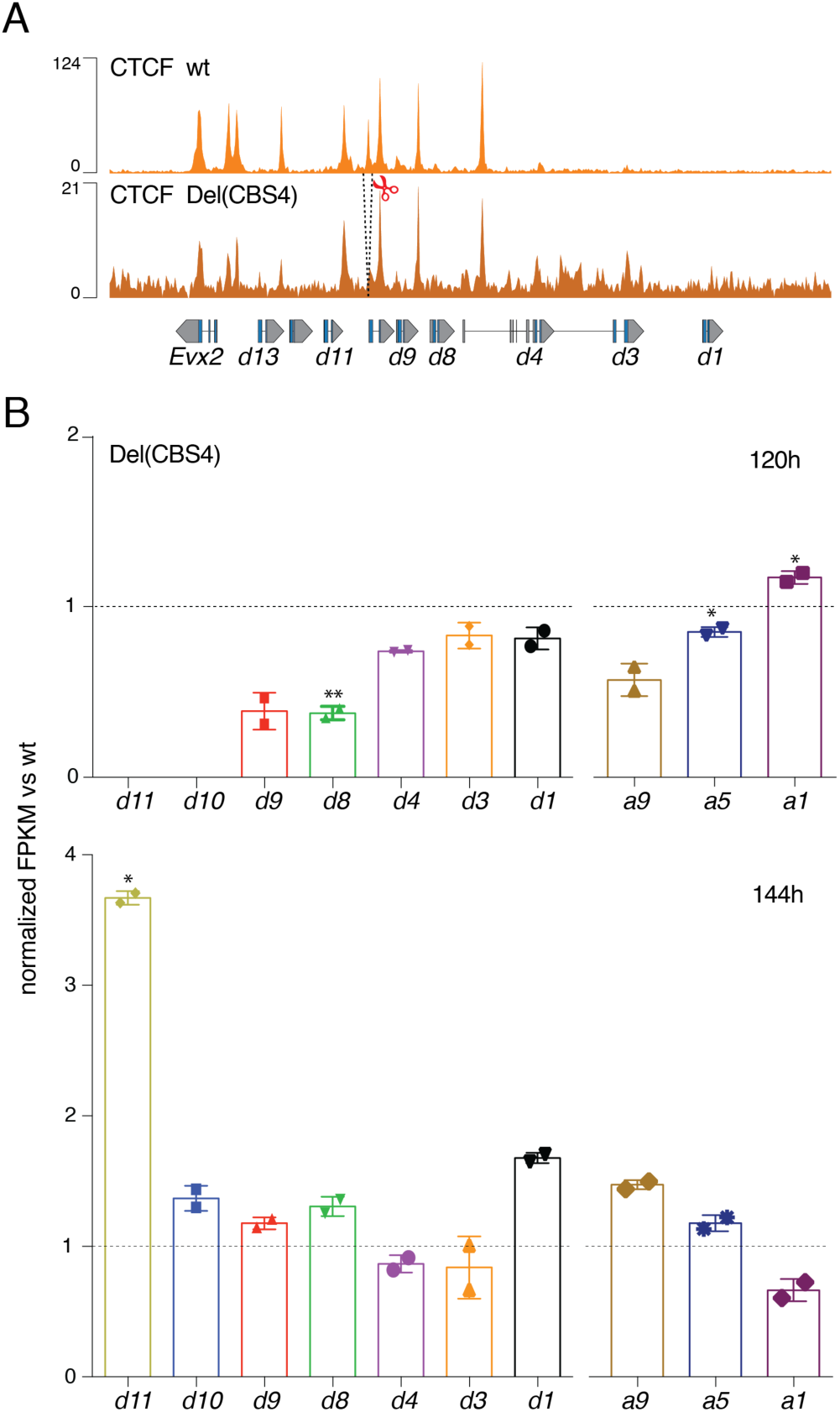
Deletion of CBS4 and effect on the surrounding *Hoxd* genes. (A) CTCF ChIP-seq of control at 168h (top) and Del(CBS4) at 120h stembryos (bottom). The deleted fragment containing CBS4 is indicated with scissors and dashed lines. (B) Normalized FPKM values of *Hoxd* and *Hoxa* genes obtained from RNA-seq libraries (n=2) produced from mutant Del(CBS4) stembryos and compared to controls at both 120h and 144h (expression of *Hoxd10* and *Hoxd11* was not considered at 120h). Values are represented as means ± SD. P-values were determined by Welch’s unequal variances t-test (* is p-value < 0.05, ** < 0.01).

**Supplementary Figure 11.**
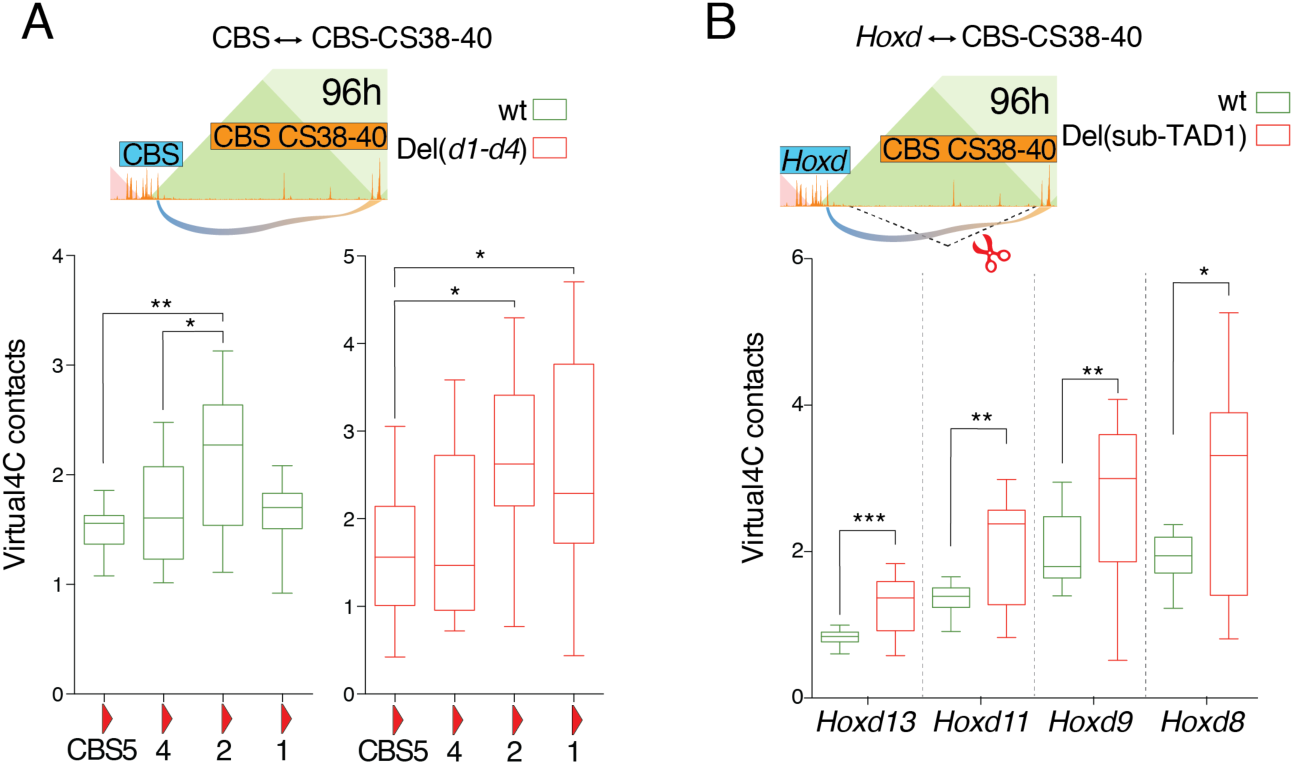
Virtual 4C contacts generated from Capture Hi-C datasets produced from either control or mutant stembryos. (A) Del(*Hoxd1-Hoxd4*) and (B) Del(sub-TAD1) stembryos were used at 96h. Data were mapped to corresponding *in-silico* reconstructed mutant genomes. (A) Quantification of the contacts between the indicated *HoxD* cluster CBS1 to 5 (viewpoint) and the CBS of the CS38-40 region. While CBS2 is the CBS the more used in wt, in Del(CBS1-4) both CBS1 and 2 are preponderant. (B) Contacts between the *Hoxd* gene bodies indicated below (viewpoints) and the CBS of the CS38-40 region. On top, scheme of the deleted fragment (dashed lines with scissors). An increase in contact is observed between *Hoxd* genes and the CS38-40 region in the mutant Del(sub-TAD1). In (*A*), asterisks represent significant changes as determined by the Kruskal-Wallis with Dunn’s multiple comparisons test. In (*B*), the Mann-Whitney nonparametric test was used to assess significant changes between wt and Del(sub-TAD1) mutant (* is p-value < 0.05, ** < 0.01, *** < 0.001).

**Supplementary Figure 12.**
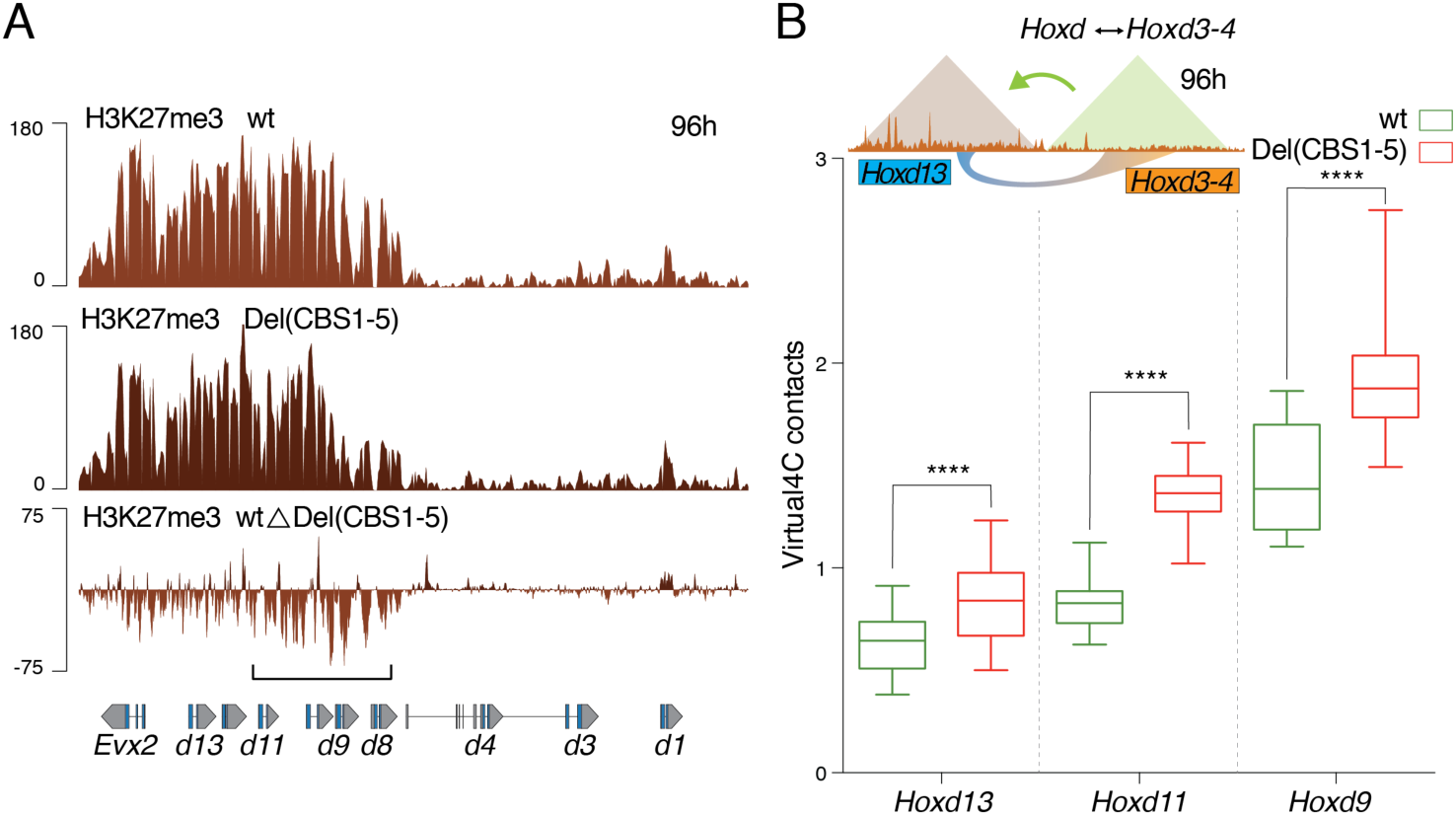
H3K27me3 profile and virtual 4C using Del(CBS1-5) stembryos. (A) H3K27me3 ChIP-seq using control (top, light brown) and Del(CBS1-5) (middle, dark brown) stembryos at 96h (average of two replicates). The subtraction track is shown below and illustrates the decrease in H3K27me3 marks over the central and posterior part of the *HoxD* cluster (bracket). (B) Virtual4C contacts generated from Capture Hi-C datasets produced out of control and Del(CBS1-5) mutant stembryos at 96h. Interactions are quantified between those *Hoxd* genes shown below (viewpoint) and the activated *Hoxd3*-*Hoxd4* part of the cluster. Increased contacts are scored in mutant stembryos for all three viewpoints, illustrating the loss, in CTCF mutant stembryos, of a clear partition of the gene cluster into two separate positive and negative domains. Asterisks represent significant changes determined by the Mann-Whitney nonparametric test (**** is p-value < 0.0001).

**Supplementary Figure 13.**
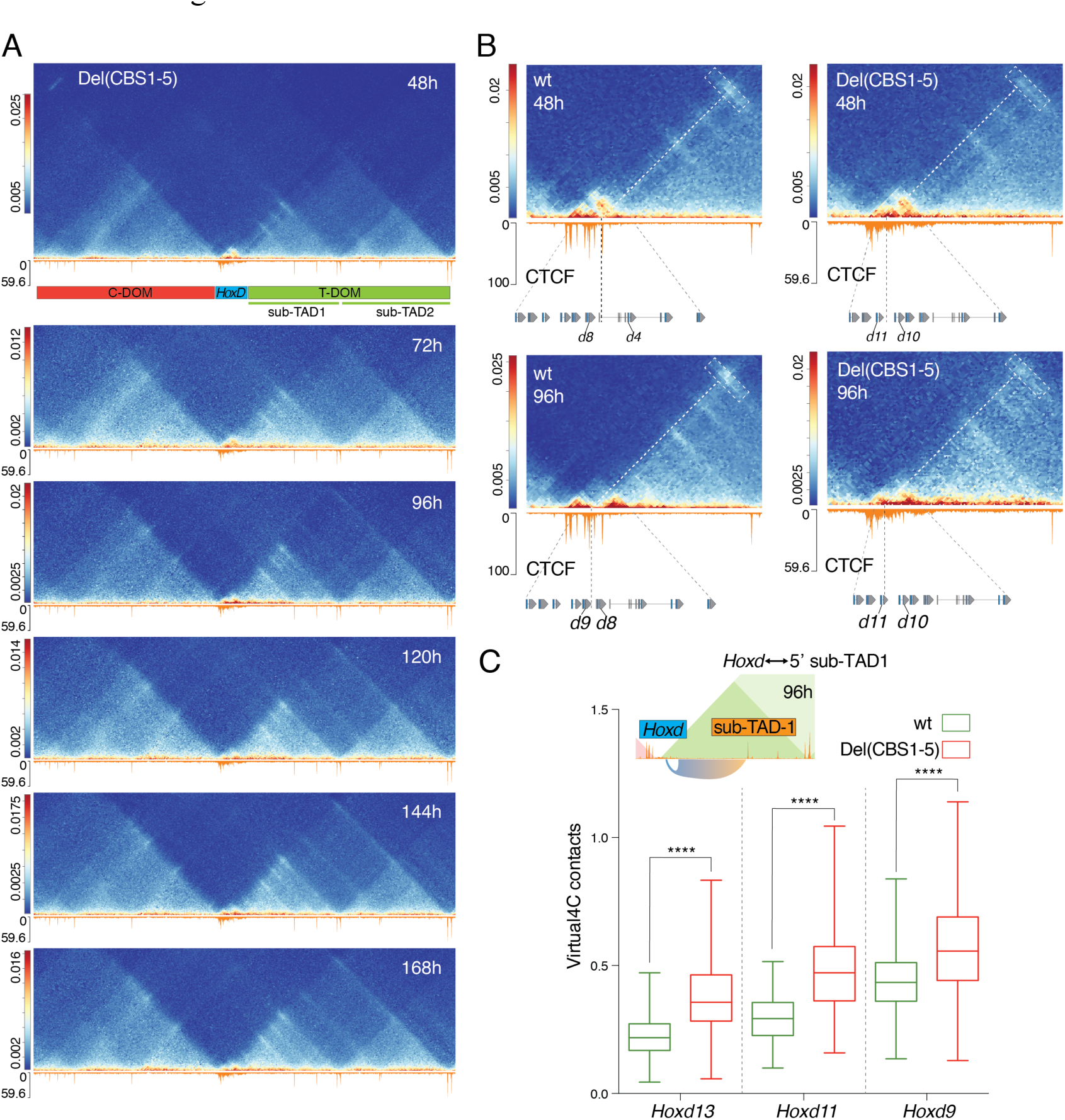
Time course capture Hi-C using Del(CBS1-5) stembryos. (A) Six time points are shown with bin size 5 kb. Libraries were mapped on a control genome (mm10). (B) Magnifications of the interactions between the *Hoxd* genes cluster and the CS38-40 region (dashed box) in control (left) and Del(CBS1-5) mutant specimen (right), either before (48h, top) or after (96h, bottom) the translocation of the anterior part of the cluster. Diagonal and vertical dashed lines indicate the position of the most prominant interactions within the *Hoxd* gene cluster with the CS38-40 region. While a shift toward a posterior position is observed between 48h and 96h in control stembryos (from CBS1 to the *Hoxd8*-*Hoxd9* interval), interactions scored in the Del(CBS1-5) mutant condition remain diffused over the gene cluster and only a slight change in the posterior limit is observed (around *Hoxd11*). (C) Virtual4C contacts generated from Capture Hi-C datasets produced in control and Del(CBS1-5) mutant stembryos at 96h. Interactions between the indicated *Hoxd* genes (viewpoints) and the 5’ region of sub-TAD1 are significantly higher in Del(CBS1-5) mutant. Asterisks represent significant changes determined by the Mann-Whitney nonparametric test (**** is p-value < 0.0001).

### Supplementary Tables 1 to 3

**Supplementary Table 1:**
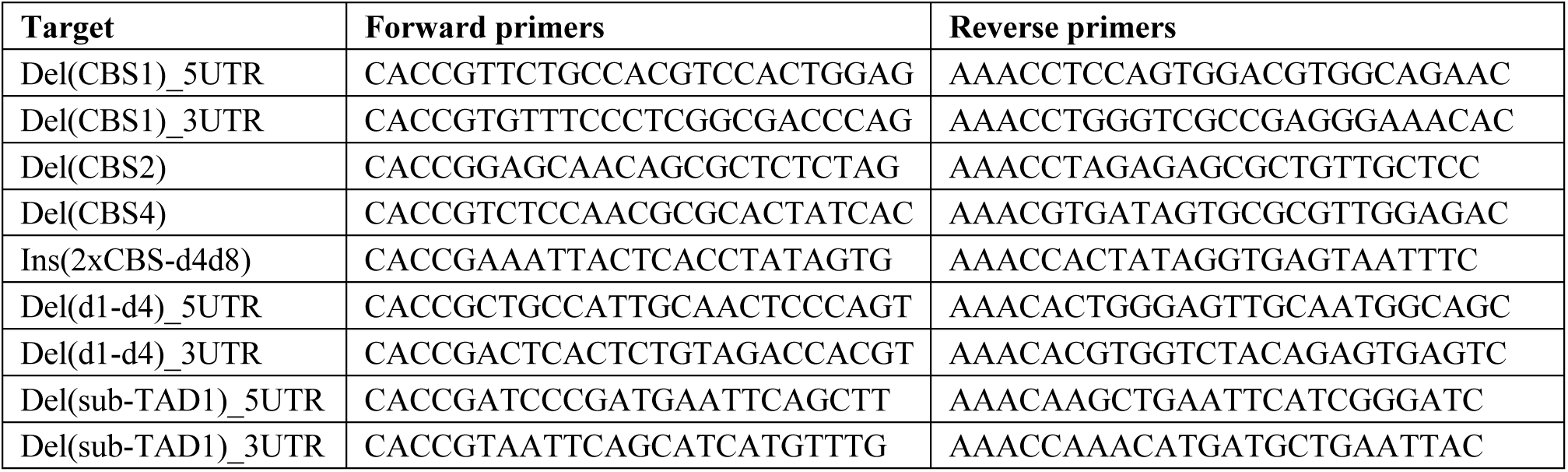
List of sgRNAs used to generate the various mutant ES cell lines as well as the Del(sub-TAD1) mutant mice.

**Supplementary Table 2:**
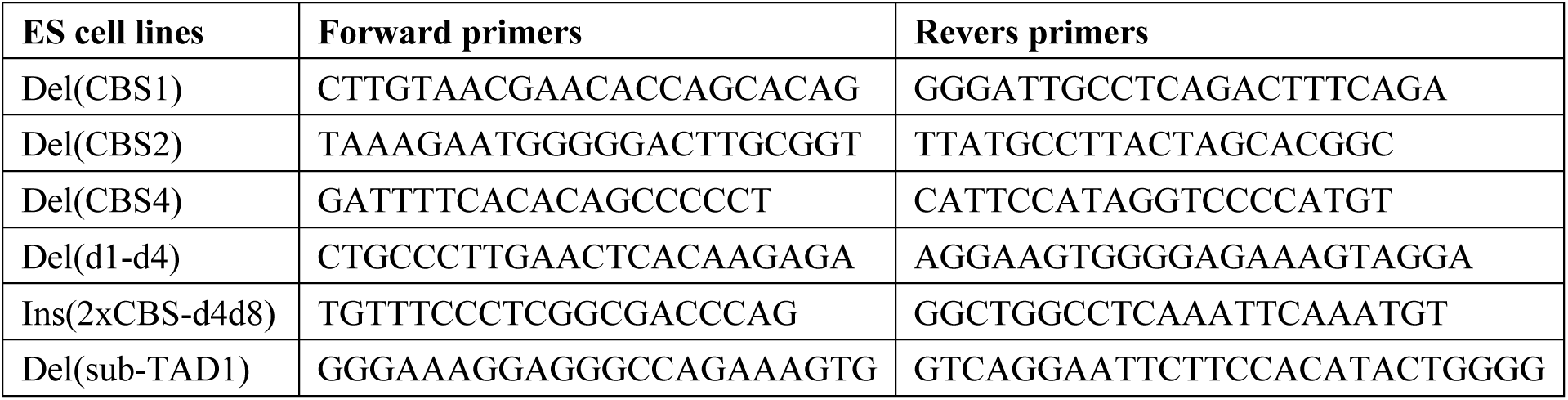
List of genotyping primers used to characterize the various mutations produced in ES cells within the *HoxD* locus.

**Supplementary Table 3:**
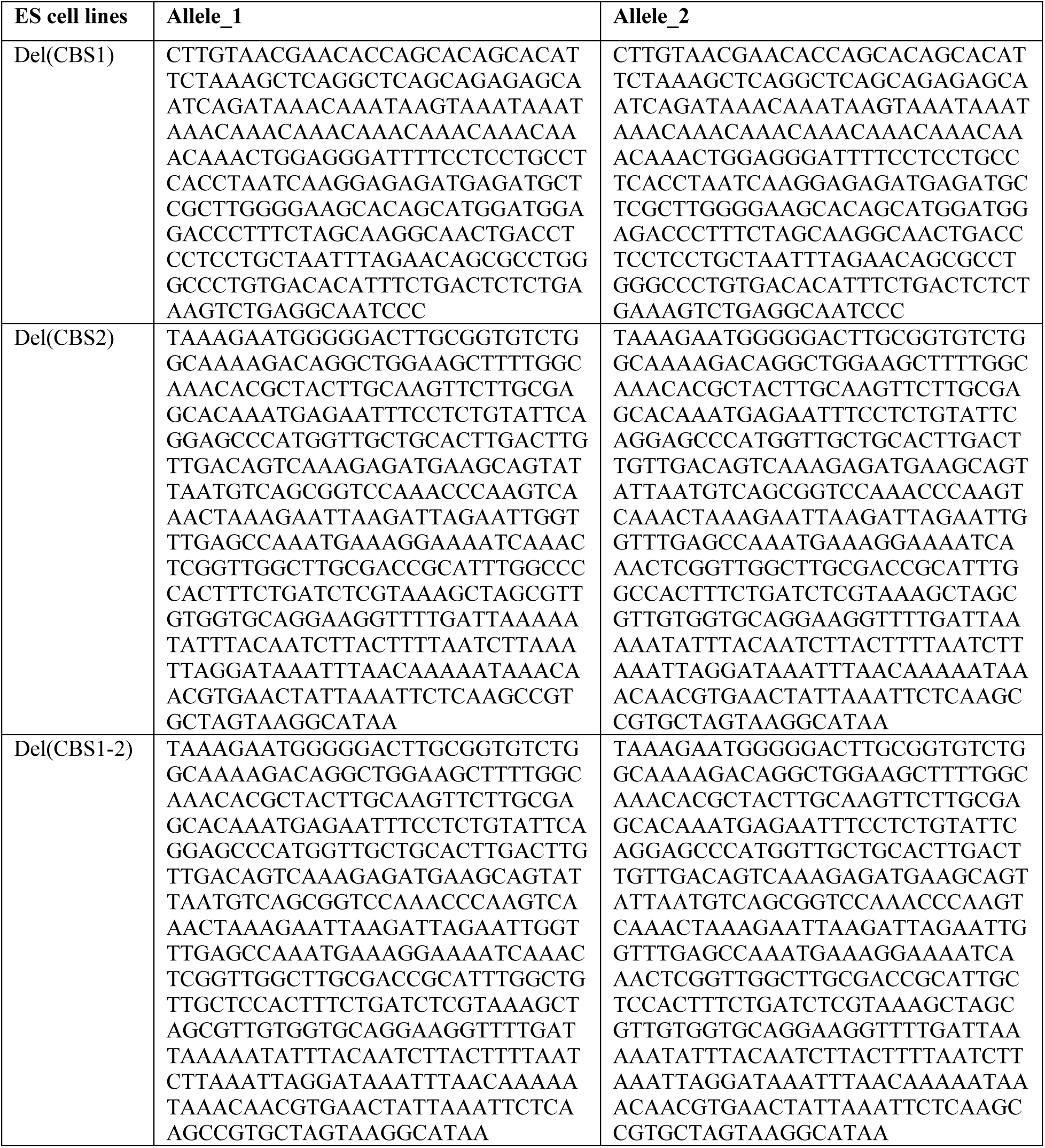
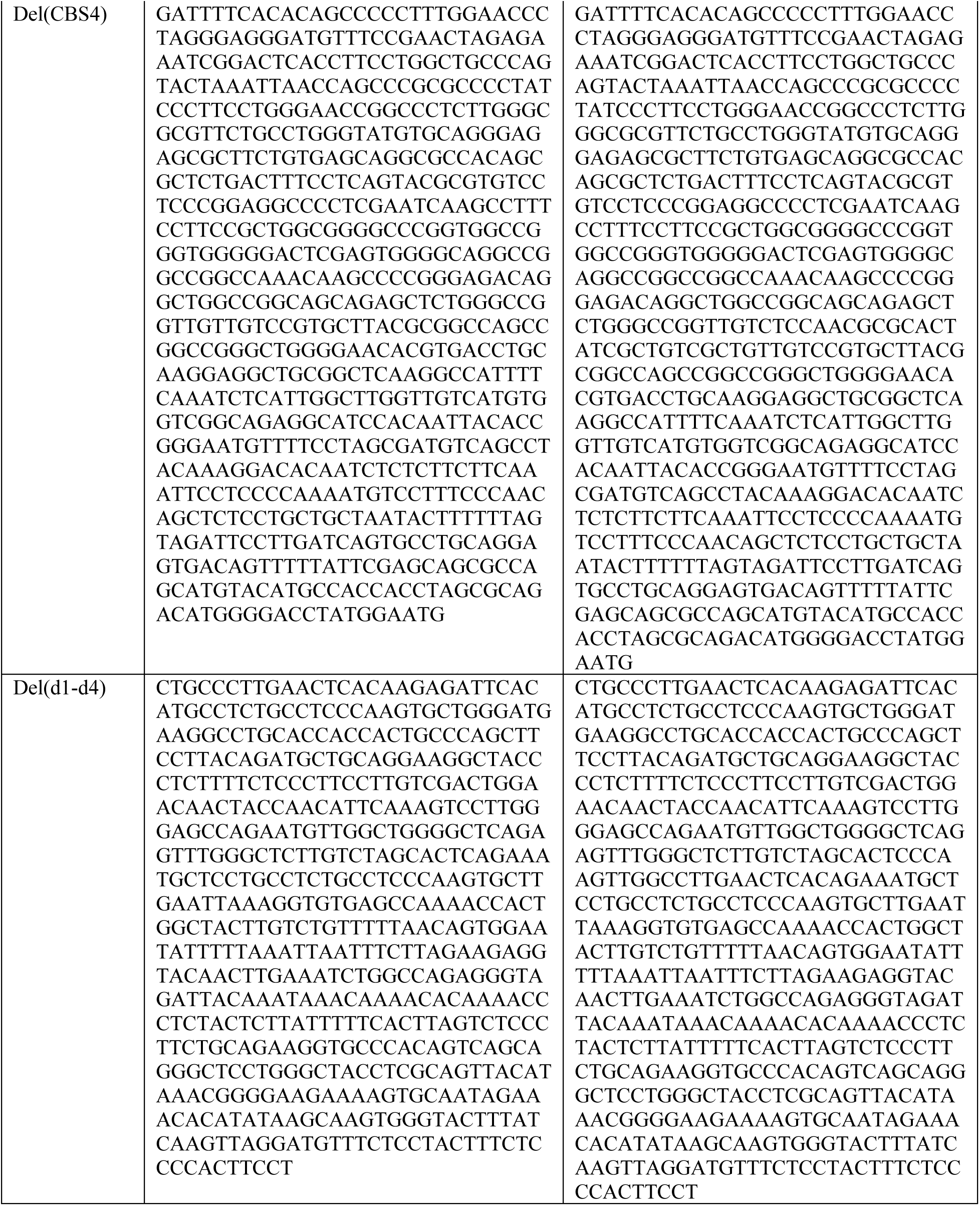
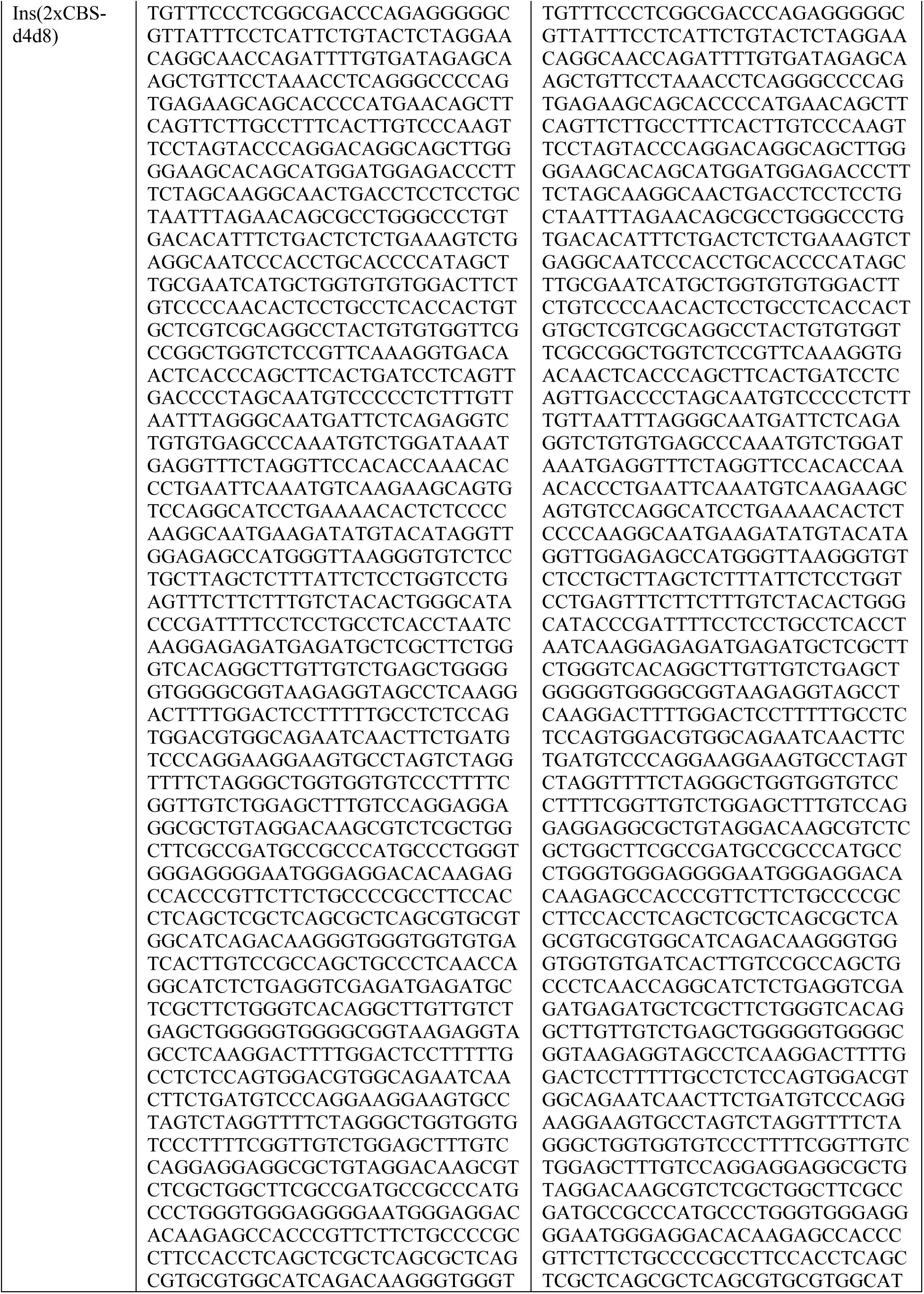
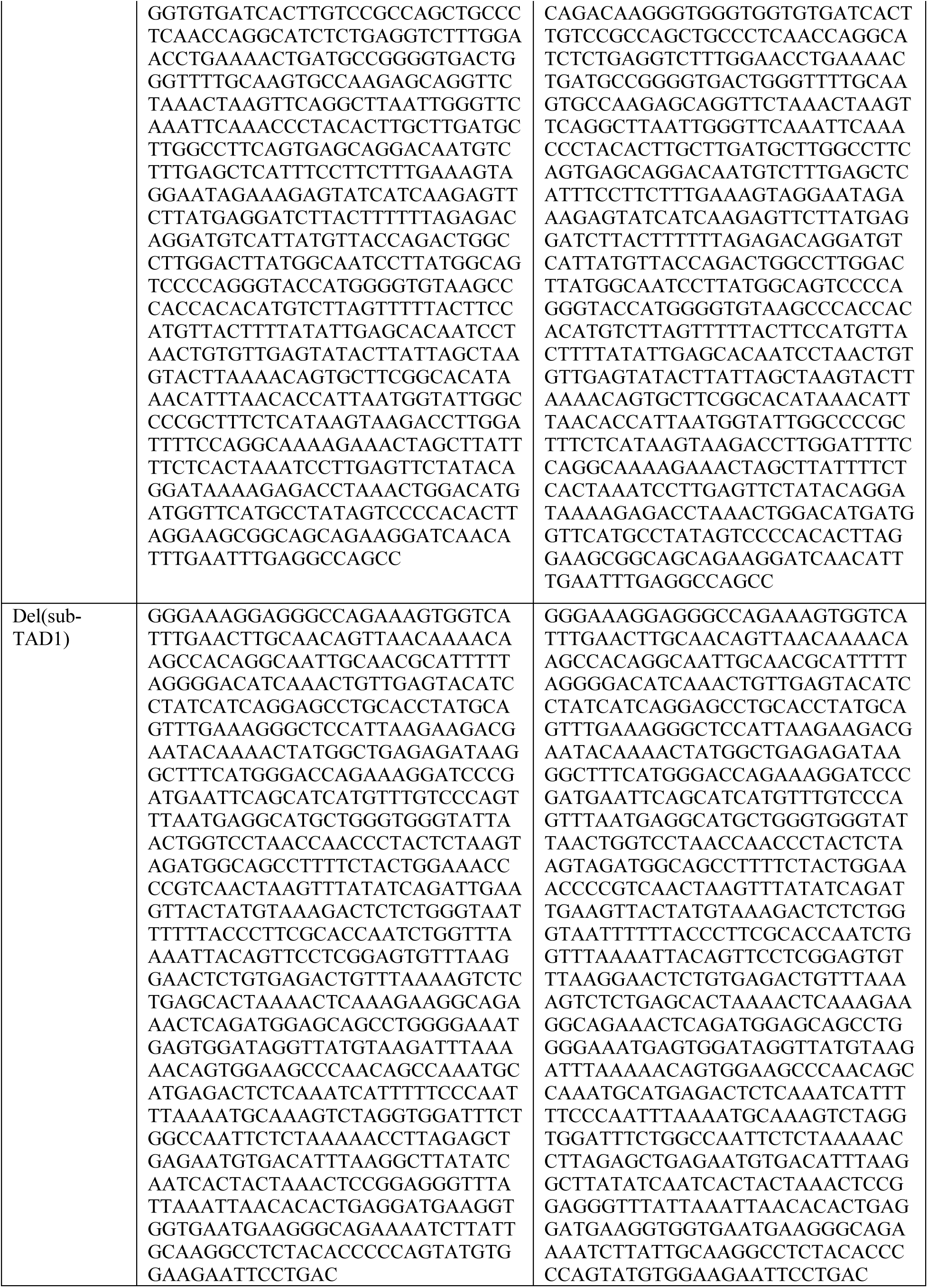
Sanger sequencing of mutant ES cell lines produced in this study. DNA sequence was reconstructed for both alleles, using the genotyping primers shown in supplementary Table S2.

### Supplementary Movies 1-2

**Supplementary Movie 1.**

https://pegasus.epfl.ch/?94641ba4de6c3fd2#78LhibFdGQQJgoik1RrmmqivLG5JedeECEbJLSRHJxXy

**Legend to Supplementary movie 1: Dynamic of colinear activation at the *HoxD* cluster.** Time-lapse reconstruction of H3K27ac coverage at the *HoxD* cluster between 72h and 168h, with a CTCF ChIP-seq profile (top). Each frame represents 1 h time resolution. A rapid coverage of H3K27ac is detected over the *Hoxd1* to *Hoxd4* region, followed by a slow and progressive spreading throughout the central and posterior part of the cluster.

Supplementary Movie 2.

https://pegasus.epfl.ch/?345f5c9c1b2c22f5#JDXq3kJ4XxdPf1f53yytizn3kJkfPRS9M17SghEbJRkA

**Legend to Supplementary movie 2: Time-lapse reconstruction of the dynamic topology at the *HoxD* locus upon transcriptional activation.** Contacts generated from the CHi-C time-course with a 1 h per frame resolution. Within the *HoxD* cluster, the initial inactive state is visible by dense intra-cluster interactions (brown arrow). The cluster then translocates and two micro-TADs appear separated by CBS1 and corresponding to the inactive posterior region (brown arrow) and the newly activated anterior region (blue arrow). Constitutive interactions between the gene cluster and T-DOM are visible at 48h with a predominance of CBS1 (bold white line). Soon after gene activation and the concurrent translocation of the cluster, this preferential interaction slowly shifts towards more central positions within the cluster (white line).

## REFERENCES

Abdennur, N., and Mirny, L.A. (2020). Cooler: scalable storage for Hi-C data and other genomically labeled arrays. Bioinformatics 36, 311–316. https://doi.org/10.1093/bioinformatics/btz540.

Afgan, E., Baker, D., van den Beek, M., Blankenberg, D., Bouvier, D., Čech, M., Chilton, J., Clements, D., Coraor, N., Eberhard, C., et al. (2016). The Galaxy platform for accessible, reproducible and collaborative biomedical analyses: 2016 update. Nucleic Acids Res. 44, W3–W10. https://doi.org/10.1093/nar/gkw343.

Aires, R., de Lemos, L., Nóvoa, A., Jurberg, A.D., Mascrez, B., Duboule, D., and Mallo, M. (2019). Tail Bud Progenitor Activity Relies on a Network Comprising Gdf11, Lin28, and Hox13 Genes. Dev. Cell 48, 383–395.e8. https://doi.org/10.1016/j.devcel.2018.12.004.

Allais-Bonnet, A., Grohs, C., Medugorac, I., Krebs, S., Djari, A., Graf, A., Fritz, S., Seichter, D., Baur, A., Russ, I., et al. (2013). Novel Insights into the Bovine Polled Phenotype and Horn Ontogenesis in Bovidae. PLoS ONE 8, e63512. https://doi.org/10.1371/journal.pone.0063512.

Amândio, A.R., Beccari, L., Lopez-Delisle, L., Mascrez, B., Zakany, J., Gitto, S., and Duboule, D. (2021). Sequential in *cis* mutagenesis in vivo reveals various functions for CTCF sites at the mouse *HoxD* cluster. Genes Dev. genesdev;gad.348934.121v1. https://doi.org/10.1101/gad.348934.121.

Amin, S., Neijts, R., Simmini, S., van Rooijen, C., Tan, S.C., Kester, L., van Oudenaarden, A., Creyghton, M.P., and Deschamps, J. (2016). Cdx and T Brachyury Co-activate Growth Signaling in the Embryonic Axial Progenitor Niche. Cell Rep 17, 3165–3177. https://doi.org/10.1016/j.celrep.2016.11.069.

Anania, C., Acemel, R.D., Jedamzick, J., Bolondi, A., Cova, G., Brieske, N., Kühn, R., Wittler, L., Real, F.M., and Lupiáñez, D.G. (2021). In vivo dissection of a clustered-CTCF domain boundary reveals developmental principles of regulatory insulation. BioRxiv 2021.04.14.439779. https://doi.org/10.1101/2021.04.14.439779.

Barnett, D.W., Garrison, E.K., Quinlan, A.R., Stromberg, M.P., and Marth, G.T. (2011). BamTools: a C++ API and toolkit for analyzing and managing BAM files. Bioinformatics 27, 1691–1692. https://doi.org/10.1093/bioinformatics/btr174.

Beccari, L., Moris, N., Girgin, M., Turner, D.A., Baillie-Johnson, P., Cossy, A.-C., Lutolf, M.P., Duboule, D., and Arias, A.M. (2018). Multi-axial self-organization properties of mouse embryonic stem cells into gastruloids. Nature 562, 272–276. https://doi.org/10.1038/s41586-018-0578-0.

Bernstein, B.E., Kamal, M., Lindblad-Toh, K., Bekiranov, S., Bailey, D.K., Huebert, D.J., McMahon, S., Karlsson, E.K., Kulbokas, E.J., Gingeras, T.R., et al. (2005). Genomic maps and comparative analysis of histone modifications in human and mouse. Cell 120, 169–181. https://doi.org/10.1016/j.cell.2005.01.001.

Bolt, C.C., Lopez-Delisle, L., Mascrez, B., and Duboule, D. (2021). Mesomelic dysplasias associated with the HOXD locus are caused by regulatory reallocations. Nat Commun 12, 5013. https://doi.org/10.1038/s41467-021-25330-y.

van den Brink, S.C., Baillie-Johnson, P., Balayo, T., Hadjantonakis, A.K., Nowotschin, S., Turner, D.A., and Martinez Arias, A. (2014). Symmetry breaking, germ layer specification and axial organisation in aggregates of mouse embryonic stem cells. Development 141, 4231–4242. https://doi.org/10.1242/dev.113001.

van den Brink, S.C., Alemany, A., van Batenburg, V., Moris, N., Blotenburg, M., Vivié, J., Baillie-Johnson, P., Nichols, J., Sonnen, K.F., Martinez Arias, A., et al. (2020). Single-cell and spatial transcriptomics reveal somitogenesis in gastruloids. Nature https://doi.org/10.1038/s41586-020-2024-3.

Burke, A.C., Nelson, C.E., Morgan, B.A., and Tabin, C. (1995). Hox genes and the evolution of vertebrate axial morphology. Development 121, 333–346. .

Busslinger, G.A., Stocsits, R.R., van der Lelij, P., Axelsson, E., Tedeschi, A., Galjart, N., and Peters, J.-M. (2017). Cohesin is positioned in mammalian genomes by transcription, CTCF and Wapl. Nature 544, 503–507. https://doi.org/10.1038/nature22063.

Ciosk, R., Shirayama, M., Shevchenko, A., Tanaka, T., Toth, A., Shevchenko, A., and Nasmyth, K. (2000). Cohesin’s Binding to Chromosomes Depends on a Separate Complex Consisting of Scc2 and Scc4 Proteins. Molecular Cell 5, 243–254. https://doi.org/10.1016/S1097-2765(00)80420-7.

Cooke, J., and Zeeman, E.C. (1976). A clock and wavefront model for control of the number of repeated structures during animal morphogenesis. Journal of Theoretical Biology 58, 455–476. https://doi.org/10.1016/S0022-5193(76)80131-2.

Darbellay, F., Bochaton, C., Lopez-Delisle, L., Mascrez, B., Tschopp, P., Delpretti, S., Zakany, J., and Duboule, D. (2019). The constrained architecture of mammalian Hox gene clusters. Proc. Natl. Acad. Sci. U.S.A. 116, 13424– 13433. https://doi.org/10.1073/pnas.1904602116.

Deschamps, J., and Duboule, D. (2017). Embryonic timing, axial stem cells, chromatin dynamics, and the Hox clock. Genes Dev 31, 1406–1416. https://doi.org/10.1101/gad.303123.117.

Despang, A., Schöpflin, R., Franke, M., Ali, S., Jerković, I., Paliou, C., Chan, W.-L., Timmermann, B., Wittler, L., Vingron, M., et al. (2019). Functional dissection of the Sox9–Kcnj2 locus identifies nonessential and instructive roles of TAD architecture. Nat Genet 51, 1263–1271. https://doi.org/10.1038/s41588-019-0466-z.

Dobin, A., Davis, C.A., Schlesinger, F., Drenkow, J., Zaleski, C., Jha, S., Batut, P., Chaisson, M., and Gingeras, T.R. (2013). STAR: ultrafast universal RNA-seq aligner. Bioinformatics 29, 15–21. https://doi.org/10.1093/bioinformatics/bts635.

Dolle, P., Izpisua-Belmonte, J.C., Falkenstein, H., Renucci, A., and Duboule, D. (1989). Coordinate expression of the murine Hox-5 complex homoeobox-containing genes during limb pattern formation. Nature 342, 767–772. https://doi.org/10.1038/342767a0.

Duboule, D. (1994). Temporal colinearity and the phylotypic progression: A basis for the stability of a vertebrate Bauplan and the evolution of morphologies through heterochrony. Development 135–142. .

Duboule, D. (2003). Time for Chronomics? Science 301, 277–277. https://doi.org/10.1126/science.301.5631.277.

Duboule, D., and Dolle, P. (1989). The Structural and Functional-Organization of the Murine Hox Gene Family Resembles That of Drosophila Homeotic Genes. Embo J 8, 1497–1505.

Durston, A.J. (2019). Vertebrate hox temporal collinearity: does it exist and what is it’s function? Cell Cycle 18, 523–530. https://doi.org/10.1080/15384101.2019.1577652.

Durston, A., Wacker, S., Bardine, N., and Jansen, H. (2012). Time space translation: a hox mechanism for vertebrate a-p patterning. Curr Genomics 13, 300–307. https://doi.org/10.2174/138920212800793375.

Ebisuya, M., and Briscoe, J. (2018). What does time mean in development? Development 145, dev164368. https://doi.org/10.1242/dev.164368.

Fudenberg, G., Imakaev, M., Lu, C., Goloborodko, A., Abdennur, N., and Mirny, L.A. (2016). Formation of Chromosomal Domains by Loop Extrusion. Cell Rep 15, 2038–2049. https://doi.org/10.1016/j.celrep.2016.04.085.

Gabriele, M., Brandão, H.B., Grosse-Holz, S., Jha, A., Dailey, G.M., Cattoglio, C., Hsieh, T.-H.S., Mirny, L., Zechner, C., and Hansen, A.S. (2022). Dynamics of CTCF- and cohesin-mediated chromatin looping revealed by live-cell imaging. Science 376, 496–501. https://doi.org/10.1126/science.abn6583.

Garcia-Fernàndez, J., and Holland, P.W.H. (1994). Archetypal organization of the amphioxus Hox gene cluster. Nature 370, 563–566. https://doi.org/10.1038/370563a0.

Gaunt, S.J. (1994). Conservation in the Hox code during morphological evolution. The International Journal of Developmental Biology 38, 549–552. .

Gaunt, S.J. (2015). The significance of Hox gene collinearity. Int J Dev Biol 59, 159–170. https://doi.org/10.1387/ijdb.150223sg.

Gaunt, S.J. (2018). Made in the Image of a Fly.

Gaunt, S., Sharpe, P.T., and Duboule, D. (1988). Spatially restricted domains of homeo-gene transcripts in mouse embryos: relation to a segmented body plan. Development 104 (*suppl*), 169–179. .

Gaunt, S.J., George, M., and Paul, Y.-L. (2013). Direct activation of a mouse Hoxd11 axial expression enhancer by Gdf11/Smad signalling. Dev Biol 383, 52–60. https://doi.org/10.1016/j.ydbio.2013.08.025.

Goldbeter, A., and Pourquié, O. (2008). Modeling the segmentation clock as a network of coupled oscillations in the Notch, Wnt and FGF signaling pathways. J Theor Biol 252, 574–585. https://doi.org/10.1016/j.jtbi.2008.01.006.

Gould, S.J. (1977). Ontogeny and Phylogeny (Belknap Press of Harvad University Press).

Graham, A., Papalopulu, N., and Krumlauf, R. (1989). The murine and Drosophila homeobox gene complexes have common features of organization and expression. Cell 57, 367–378. .

Harding, K., Wedeen, C., McGinnis, W., and Levine, M. (1985). Spatially regulated expression of homeotic genes in Drosophila. Science 229, 1236–1242. .

Izpisua-Belmonte, J.C., Falkenstein, H., Dolle, P., Renucci, A., and Duboule, D. (1991). Murine genes related to the Drosophila AbdB homeotic genes are sequentially expressed during development of the posterior part of the body. The EMBO Journal 10, 2279–2289. .

Kagey, M.H., Newman, J.J., Bilodeau, S., Zhan, Y., Orlando, D.A., van Berkum, N.L., Ebmeier, C.C., Goossens, J., Rahl, P.B., Levine, S.S., et al. (2010). Mediator and cohesin connect gene expression and chromatin architecture. Nature 467, 430–435. https://doi.org/10.1038/nature09380.

Kessel, M., and Gruss, P. (1991). Homeotic transformations of murine vertebrae and concomitant alteration of Hox codes induced by retinoic acid. Cell 67, 89–104. .

Kmita, M., and Duboule, D. (2003). Organizing axes in time and space; 25 years of colinear tinkering. Science 301, 331–333. https://doi.org/10.1126/science.1085753.

Kondo, M., Matsuo, M., Igarashi, K., Haramoto, Y., Yamamoto, T., Yasuoka, Y., and Taira, M. (2019). *de novo* transcription of multiple Hox cluster genes takes place simultaneously in early *Xenopus tropicalis* embryos. Biology Open bio.038422. https://doi.org/10.1242/bio.038422.

Kraft, K., Magg, A., Heinrich, V., Riemenschneider, C., Schöpflin, R., Markowski, J., Ibrahim, D.M., Acuna-Hidalgo, R., Despang, A., Andrey, G., et al. (2019). Serial genomic inversions induce tissue-specific architectural stripes, gene misexpression and congenital malformations. Nat Cell Biol 21, 305–310. https://doi.org/10.1038/s41556-019-0273-x.

Krumlauf, R. (1994). Hox genes in vertebrate development. Cell 78, 191–201. .

Langmead, B., and Salzberg, S.L. (2012). Fast gapped-read alignment with Bowtie 2. Nat Methods 9, 357–359. https://doi.org/10.1038/nmeth.1923.

Lewis, E.B. (1978). A gene complex controlling segmentation in Drosophila. Nature 276, 565–570. .

Lopez-Delisle, L., Rabbani, L., Wolff, J., Bhardwaj, V., Backofen, R., Grüning, B., Ramírez, F., and Manke, T. (2021). pyGenomeTracks: reproducible plots for multivariate genomic datasets. Bioinformatics 37, 422–423. https://doi.org/10.1093/bioinformatics/btaa692.

Lopez-Delisle, Lucille (2021). Customized gtf file from Ensembl version 102 mm10 (Zenodo).

Martin, M. (2011). Cutadapt removes adapter sequences from high-throughput sequencing reads. EMBnet.Journal 17. https://doi.org/10.14806/ej.17.1.200.

Mazzoni, E.O., Mahony, S., Peljto, M., Patel, T., Thornton, S.R., McCuine, S., Reeder, C., Boyer, L.A., Young, R.A., Gifford, D.K., et al. (2013). Saltatory remodeling of Hox chromatin in response to rostrocaudal patterning signals. Nat Neurosci 16, 1191–1198. https://doi.org/10.1038/nn.3490.

Mumbach, M.R., Rubin, A.J., Flynn, R.A., Dai, C., Khavari, P.A., Greenleaf, W.J., and Chang, H.Y. (2016). HiChIP: efficient and sensitive analysis of protein-directed genome architecture. Nat Methods 13, 919–922. https://doi.org/10.1038/nmeth.3999.

Narendra, V., Rocha, P.P., An, D., Raviram, R., Skok, J.A., Mazzoni, E.O., and Reinberg, D. (2015). CTCF establishes discrete functional chromatin domains at the Hox clusters during differentiation. Science 347, 1017– 1021. https://doi.org/10.1126/science.1262088.

Narendra, V., Bulajić, M., Dekker, J., Mazzoni, E.O., and Reinberg, D. (2016). CTCF-mediated topological boundaries during development foster appropriate gene regulation. Genes Dev. 30, 2657–2662. https://doi.org/10.1101/gad.288324.116.

Neijts, R., and Deschamps, J. (2017). At the base of colinear Hox gene expression: cis -features and trans -factors orchestrating the initial phase of Hox cluster activation. Developmental Biology 428, 293–299. https://doi.org/10.1016/j.ydbio.2017.02.009.

Neijts, R., Amin, S., van Rooijen, C., Tan, S., Creyghton, M.P., de Laat, W., and Deschamps, J. (2016). Polarized regulatory landscape and Wnt responsiveness underlie Hox activation in embryos. Genes Dev 30, 1937–1942. https://doi.org/10.1101/gad.285767.116.

Neijts, R., Amin, S., van Rooijen, C., and Deschamps, J. (2017). Cdx is crucial for the timing mechanism driving colinear Hox activation and defines a trunk segment in the Hox cluster topology. Dev Biol 422, 146–154. https://doi.org/10.1016/j.ydbio.2016.12.024.

Noordermeer, D., Leleu, M., Splinter, E., Rougemont, J., De Laat, W., and Duboule, D. (2011). The dynamic architecture of Hox gene clusters. Science 334, 222–225. https://doi.org/10.1126/science.1207194.

Noordermeer, D., Leleu, M., Schorderet, P., Joye, E., Chabaud, F., and Duboule, D. (2014). Temporal dynamics and developmental memory of 3D chromatin architecture at Hox gene loci. ELife 3, e02557. https://doi.org/10.7554/eLife.02557.

Ortabozkoyun, H., Huang, P.-Y., Cho, H., Narendra, V., LeRoy, G., Gonzalez-Buendia, E., Skok, J.A., Tsirigos, A., Mazzoni, E.O., and Reinberg, D. (2022). CRISPR and biochemical screens identify MAZ as a cofactor in CTCF-mediated insulation at Hox clusters. Nat Genet 54, 202–212. https://doi.org/10.1038/s41588-021-01008-5.

Palmeirim, I., Henrique, D., Ish-Horowicz, D., and Pourquie, O. (1997). Avian hairy gene expression identifies a molecular clock linked to vertebrate segmentation and somitogenesis. Cell 91, 639–648. .

Ramirez, F., Ryan, D.P., Gruning, B., Bhardwaj, V., Kilpert, F., Richter, A.S., Heyne, S., Dundar, F., and Manke, T. (2016). deepTools2: a next generation web server for deep-sequencing data analysis. Nucleic Acids Research 44, W160–5. https://doi.org/10.1093/nar/gkw257.

Rekaik, H., and Lopez-Delisle, L. (2022). chr2 of mutant genomes used in Rekaik et al. 2022 (Zenodo).

Roberts, A., Trapnell, C., Donaghey, J., Rinn, J.L., and Pachter, L. (2011). Improving RNA-Seq expression estimates by correcting for fragment bias. Genome Biology 12, R22. https://doi.org/10.1186/gb-2011-12-3-r22.

Rodriguez-Carballo, E., Lopez-Delisle, L., Zhan, Y., Fabre, P.J., Beccari, L., El-Idrissi, I., Huynh, T.H.N., Ozadam, H., Dekker, J., and Duboule, D. (2017). The HoxD cluster is a dynamic and resilient TAD boundary controlling the segregation of antagonistic regulatory landscapes. Genes Dev. 31, 2264–2281. https://doi.org/10.1101/gad.307769.117.

Rodríguez-Carballo, E., Lopez-Delisle, L., Willemin, A., Beccari, L., Gitto, S., Mascrez, B., and Duboule, D. (2020). Chromatin topology and the timing of enhancer function at the *HoxD* locus. Proc Natl Acad Sci USA 117, 31231–31241. https://doi.org/10.1073/pnas.2015083117.

Sanborn, A.L., Rao, S.S.P., Huang, S.-C., Durand, N.C., Huntley, M.H., Jewett, A.I., Bochkov, I.D., Chinnappan, D., Cutkosky, A., Li, J., et al. (2015). Chromatin extrusion explains key features of loop and domain formation in wild-type and engineered genomes. Proc. Natl. Acad. Sci. U.S.A. 112, E6456–6465. https://doi.org/10.1073/pnas.1518552112.

Silva, J., Barrandon, O., Nichols, J., Kawaguchi, J., Theunissen, T.W., and Smith, A. (2008). Promotion of Reprogramming to Ground State Pluripotency by Signal Inhibition. PLoS Biol 6, e253. https://doi.org/10.1371/journal.pbio.0060253.

Soshnikova, N., and Duboule, D. (2009). Epigenetic temporal control of mouse Hox genes in vivo. Science 324, 1320–1323. https://doi.org/10.1126/science.1171468.

Trapnell, C., Williams, B.A., Pertea, G., Mortazavi, A., Kwan, G., van Baren, M.J., Salzberg, S.L., Wold, B.J., and Pachter, L. (2010). Transcript assembly and quantification by RNA-Seq reveals unannotated transcripts and isoform switching during cell differentiation. Nature Biotechnology 28, 511–515. https://doi.org/10.1038/nbt.1621.

Turner, D.A., Girgin, M., Alonso-Crisostomo, L., Trivedi, V., Baillie-Johnson, P., Glodowski, C.R., Hayward, P.C., Collignon, J., Gustavsen, C., Serup, P., et al. (2017). Anteroposterior polarity and elongation in the absence of extraembryonic tissues and spatially localised signalling in Gastruloids, mammalian embryonic organoids. Development 144, 3894–3906. https://doi.org/10.1242/dev.150391.

Tzouanacou, E., Wegener, A., Wymeersch, F.J., Wilson, V., and Nicolas, J.F. (2009). Redefining the progression of lineage segregations during mammalian embryogenesis by clonal analysis. Dev Cell 17, 365–376. https://doi.org/10.1016/j.devcel.2009.08.002.

Veenvliet, J.V., Bolondi, A., Kretzmer, H., Haut, L., Scholze-Wittler, M., Schifferl, D., Koch, F., Guignard, L., Kumar, A.S., Pustet, M., et al. (2020). Mouse embryonic stem cells self-organize into trunk-like structures with neural tube and somites. Science 370. https://doi.org/10.1126/science.aba4937.

Veenvliet, J.V., Lenne, P.-F., Turner, D.A., Nachman, I., and Trivedi, V. (2021). Sculpting with stem cells: how models of embryo development take shape. Development 148, dev192914. https://doi.org/10.1242/dev.192914.

Vian, L., Pękowska, A., Rao, S.S.P., Kieffer-Kwon, K.-R., Jung, S., Baranello, L., Huang, S.-C., El Khattabi, L., Dose, M., Pruett, N., et al. (2018). The Energetics and Physiological Impact of Cohesin Extrusion. Cell 175, 292–294. https://doi.org/10.1016/j.cell.2018.09.002.

Wendt, K.S., Yoshida, K., Itoh, T., Bando, M., Koch, B., Schirghuber, E., Tsutsumi, S., Nagae, G., Ishihara, K., Mishiro, T., et al. (2008). Cohesin mediates transcriptional insulation by CCCTC-binding factor. Nature 451, 796– 801. https://doi.org/10.1038/nature06634.

Wilson, V., Olivera-Martinez, I., and Storey, K.G. (2009). Stem cells, signals and vertebrate body axis extension. Development 136, 1591–1604. https://doi.org/10.1242/dev.021246.

Wingett, S., Ewels, P., Furlan-Magaril, M., Nagano, T., Schoenfelder, S., Fraser, P., and Andrews, S. (2015). HiCUP: pipeline for mapping and processing Hi-C data. F1000Res 4, 1310. https://doi.org/10.12688/f1000research.7334.1.

Wutz, G., Várnai, C., Nagasaka, K., Cisneros, D.A., Stocsits, R.R., Tang, W., Schoenfelder, S., Jessberger, G., Muhar, M., Hossain, M.J., et al. (2017). Topologically associating domains and chromatin loops depend on cohesin and are regulated by CTCF, WAPL, and PDS5 proteins. EMBO J 36, 3573–3599. https://doi.org/10.15252/embj.201798004.

Yakushiji-Kaminatsui, N., Lopez-Delisle, L., Bolt, C.C., Andrey, G., Beccari, L., and Duboule, D. (2018). Similarities and differences in the regulation of HoxD genes during chick and mouse limb development. PLoS Biol. 16, e3000004. https://doi.org/10.1371/journal.pbio.3000004.

Young, T., Rowland, J.E., van de Ven, C., Bialecka, M., Novoa, A., Carapuco, M., van Nes, J., de Graaff, W., Duluc, I., Freund, J.N., et al. (2009). Cdx and Hox genes differentially regulate posterior axial growth in mammalian embryos. Developmental Cell 17, 516–526. https://doi.org/10.1016/j.devcel.2009.08.010.

Zaborowska, J., Egloff, S., and Murphy, S. (2016). The pol II CTD: new twists in the tail. Nat Struct Mol Biol 23, 771–777. https://doi.org/10.1038/nsmb.3285.

Zakany, J., Kmita, M., and Duboule, D. (2004). A dual role for Hox genes in limb anterior-posterior asymmetry. Science 304, 1669–1672. https://doi.org/10.1126/science.1096049.

Zhu, Y., Denholtz, M., Lu, H., and Murre, C. (2021). Calcium signaling instructs NIPBL recruitment at active enhancers and promoters via distinct mechanisms to reconstruct genome compartmentalization. Genes Dev 35, 65–81. https://doi.org/10.1101/gad.343475.120.

Zuin, J., Franke, V., van IJcken, W.F.J., van der Sloot, A., Krantz, I.D., van der Reijden, M.I.J.A., Nakato, R., Lenhard, B., and Wendt, K.S. (2014). A Cohesin-Independent Role for NIPBL at Promoters Provides Insights in CdLS. PLoS Genet 10, e1004153. https://doi.org/10.1371/journal.pgen.1004153.

